# Decomposing neural responses to melodic surprise in musicians and non-musicians: evidence for a hierarchy of predictions in the auditory system

**DOI:** 10.1101/786574

**Authors:** D.R. Quiroga-Martinez, N.C. Hansen, A. Højlund, M. Pearce, E. Brattico, P. Vuust

## Abstract

Neural responses to auditory surprise are typically studied with highly unexpected, disruptive sounds. Consequently, little is known about auditory prediction in everyday contexts that are characterized by fine-grained, non-disruptive fluctuations of auditory surprise. To address this issue, we used IDyOM, a computational model of auditory expectation, to obtain continuous surprise estimates for a set of newly composed melodies. Our main goal was to assess whether the neural correlates of non-disruptive surprising sounds in a musical context are affected by musical expertise. Using magnetoencephalography (MEG), auditory responses were recorded from musicians and non-musicians while they listened to the melodies. Consistent with a previous study, the amplitude of the N1m component increased with higher levels of computationally estimated surprise. This effect, however, was not different between the two groups. Further analyses offered an explanation for this finding: Pitch interval size itself, rather than probabilistic prediction, was responsible for the modulation of the N1m, thus pointing to low-level sensory adaptation as the underlying mechanism. In turn, the formation of auditory regularities and proper probabilistic prediction were reflected in later components: the mismatch negativity (MMNm) and the P3am, respectively. Overall, our findings reveal a hierarchy of expectations in the auditory system and highlight the need to properly account for sensory adaptation in research addressing statistical learning.

**Highlights:**

- In melodies, sound expectedness (modeled with IDyOM) is associated with the amplitude of the N1m.
- This effect is not different between musicians and non-musicians.
- Sensory adaptation related to melodic pitch intervals explains better the N1m effect.
- Auditory regularities and the expectations captured by IDyOM are reflected in the MMNm and P3am.
- Evidence for a hierarchy of auditory predictions during melodic listening.

## 1. Introduction

Surprising sounds in auditory sequences generate neural prediction error responses (den Ouden, Kok, & de Lange, 2012). These are thought to reflect the degree to which internal predictive models are updated by novel information (Friston, 2005; Friston, Rosch, Parr, Price, & Bowman, 2017; Lieder, Stephan, Daunizeau, Garrido, & Friston, 2013). However, most research on auditory surprise has employed very simple and repetitive stimuli, occasionally disrupted by highly unexpected deviant sounds (Heilbron & Chait, 2018). Consequently, little is known about prediction in everyday auditory environments that are characterized by fine-grained, non-disruptive changes in auditory surprise.

One potential way to address this issue is to employ computational modeling to produce continuous estimates of auditory surprise in realistic stimuli. This approach was adopted by Omigie and colleagues (2013), who used Information Dynamics of Music (IDyOM) (Pearce, 2005, 2018), a variable-order Markov model of auditory expectation, to estimate note-by-note surprise in a set of melodies. Using electroencephalography (EEG), they found that the amplitude of the N1 component of the event related potential (ERP) became larger as the estimated surprise of the tones increased. This suggests that it is possible to record neural responses to subtle changes in auditory expectedness in more realistic settings.

Importantly, IDyOM incorporates both a short-term (*stm*) component, which derives dynamic expectations from the statistics of the current stimulus, and a long-term (*ltm*) component that derives schematic expectations from a training corpus. The latter simulates the knowledge of auditory signals that a listener acquires during her life-span and therefore could be a good model of auditory enculturation (Morrison, Demorest, & Pearce, 2018). Interestingly, behavioral studies have shown that IDyOM’s surprise estimates are more strongly associated with expectedness ratings in musicians compared to non-musicians (Hansen & Pearce, 2014; Hansen, Vuust, & Pearce, 2016). This has been interpreted as an expertise-related enhancement in the accuracy of internal predictive models, thus providing support to IDyOM as a model of auditory enculturation. However, it remains unknown whether similar signatures of long-term statistical learning can also be observed in the neural responses to non-disruptive auditory surprise, as modeled with IDyOM.

In the present work, we used magnetoencephalography (MEG) to address this question by recording magnetic correlates of neural activity while musicians and non-musicians listened passively to melodic sequences. Following the results of Omigie et al. (2013), we expected larger magnetic N1 (N1m)^1^ amplitudes with increasing levels of estimated surprise. Crucially, the association between surprise and neural responses was expected to be stronger in musically trained participants because their more precise musical knowledge—we conjectured—would match better the ideal observer model entailed by IDyOM. This would provide evidence for a modulation of neural activity by auditory enculturation.

In order to gain a deeper understanding of the nature of the expectations reflected in the N1m, we conducted two sets of exploratory analyses. As will be seen, these analyses were crucial for the interpretation of the results from the comparison between musicians and non-musicians. First, we aimed to disentangle the contribution of individual components of the computational model by assessing the explanatory power of different IDyOM configurations. We contrasted configurations including short-term or long-term components; configurations predicting representations of pitch interval (the pitch distance between consecutive tones) and scale degree (the pitch interval between a tone and another tone perceived as the tonal center of the context); and configurations with different combinations of these factors (see section 3.2 for further details). These comparisons aimed to reveal, for example, the extent to which participants’ expectations relied on long-term schematic knowledge relative to short-term knowledge, or on pitch interval representations relative to scale-degree representations.

Second, we compared—in sensor and source space—the N1m modulation with the magnetic counterpart of the mismatch negativity (MMNm), which is a well-studied brain response to the violation of auditory regularities (Bendixen, SanMiguel, & Schröger, 2012; Garrido, Kilner, Stephan, & Friston, 2009; Näätänen, Gaillard, & Mäntysalo, 1978; Näätänen, Paavilainen, Rinne, & Alho, 2007). Since our dataset was recorded employing a novel MMN experimental paradigm with realistic non-repetitive melodies as stimuli (Quiroga-Martinez et al., 2019b, 2019a), it provided a valuable opportunity to compare these responses in the same subjects. This comparison aimed to determine whether the N1m modulation could be better interpreted as an MMNm, something that was not clear from the results in Omigie et al., (2013). This is interesting because, although these components have similar scalp topography and latency, the N1 is thought to reflect stimulus-specific adaptation (SSA)—a process whereby neurons become less responsive to repeated sensory stimulation—(May & Tiitinen, 2010; Näätänen & Picton, 1987; Ulanovsky, Las, & Nelken, 2003), whereas the MMN has been proposed to reflect the violation of auditory predictive models (Bendixen et al., 2012; Garrido et al., 2009). Therefore, since the type of expectations modeled by IDyOM are much closer to the ones that would give rise to the MMN—or similar responses such as the Early Right Anterior Negativity (ERAN) (Koelsch, Gunter, Friederici, & Schröger, 2000)—it would be surprising to see such an early sensory component as the N1m, but not the MMNm, being modulated. This distinction is crucial when considering musical expertise, as one would expect probabilistic prediction, rather than SSA, to be modulated by accurate knowledge of the statistical regularities of a musical style.

Overall, this study sought to unveil the nature of the expectations reflected in neural responses to non-disruptive surprise, as well as the effect of expertise on them. Anticipating the results, the lack of differences between musicians and non-musicians, the fact that pitch-interval models show the best performance, and the clear dissociation between the N1m and the MMNm, all point to a rather surprising set of conclusions: that SSA, instead of probabilistic prediction, underlies the modulation of the N1m; that pitch-interval size alone better explains this effect; and that auditory regularities and higher-order probabilistic predictions are reflected in later components such as the MMNm and the P3am.

## 2. Methods

The data, code and materials necessary to reproduce these experiments and results are openly available at: https://osf.io/my6te/; DOI: 10.17605/OSF.IO/MY6TE

### 2.1. Participants

Twenty-six musicians and 24 non-musicians took part in the experiment (see Table 1 for demographics). All participants were neurologically healthy, right-handed and did not possess absolute pitch. Musical expertise was assessed with the musical training subscale of the Goldsmiths Musical Sophistication Index (GMSI) (Müllensiefen, Gingras, Musil, & Stewart, 2014). Musical skills were measured with the rhythm and melody sections of the Musical Ear Test (MET) (Wallentin, Nielsen, Friis-Olivarius, Vuust, & Vuust, 2010). GMSI (t = 16.5, p < .001) and MET total scores (t = 5.2, p < .001) were significantly higher for musicians than for non-musicians. Moreover, most musicians played pitched instruments, the most common being the piano. See supplementary file 1 for a full report of instruments played, and supporting file 4 in Quiroga-Martinez et al. (2019a) for a detailed report of individual items of the GMSI subscale. Participants were recruited through an online database and agreed to take part in the experiment voluntarily. All participants gave informed consent and were paid 300 Danish kroner (approximately 40 euro) as compensation. Two musicians (not included in the reported demographics) were excluded from the analysis due to strong artefacts caused by dental implants. The study was approved by the Regional Ethics Committee (De Videnskabsetiske Komitéer for Region Midtjylland in Denmark) and conducted in accordance with the Helsinki Declaration.

**Table 1.**
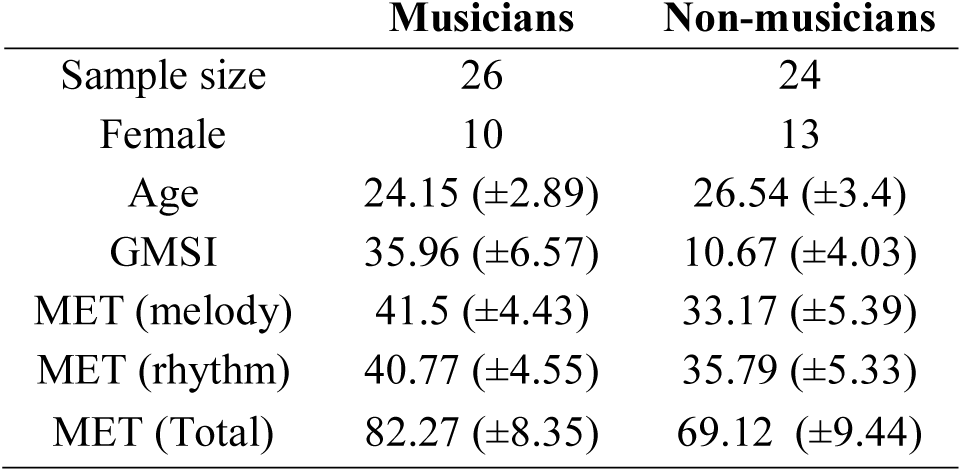
Participants’ demographic and musical expertise information. Mean and standard deviation are reported.

|  | **Musicians** | **Non-musicians** |
| --- | --- | --- |
| Sample size | 26 | 24 |
| Female | 10 | 13 |
| Age | 24.15 ($\pm 2.89$) | 26.54 ($\pm 3.4$) |
| GMSI | 35.96 ($\pm 6.57$) | 10.67 ($\pm 4.03$) |
| MET (melody) | 41.5 ($\pm 4.43$) | 33.17 ($\pm 5.39$) |
| MET (rhythm) | 40.77 ($\pm 4.55$) | 35.79 ($\pm 5.33$) |
| MET (Total) | 82.27 ($\pm 8.35$) | 69.12 ($\pm 9.44$) |

### 2.2. Stimuli

The stimuli corresponded to a set of six novel melodies composed following the rules of classical Western tonal music. These melodies were used in a recent MMN experiment (high entropy condition in Quiroga-Martinez et al. 2019b). Each melody was 32 notes long and lasted eight seconds. Major- and minor-mode versions of the melodies were transposed to six different keys (C, C#, D, D#, E, F) and were presented pseudorandomly one after the other so that no melody was repeated before all melodies were played. The major and minor versions of each melody were repeated twelve times, in randomly selected transpositions. The stimuli were delivered in three blocks lasting around seven minutes each and were part of a longer experimental session that included other conditions addressing questions beyond the scope of this study. Data from these conditions have been (Quiroga-Martinez et al. 2019a, 2019b) and will be published elsewhere. Individual tones were created using a piano sample and had a duration of 250 ms. The pitch range of the melodies, as presented during the session, spanned 31 semitones from B3 (F_0_ ≈ 247 Hz) to F6 (F_0_ ≈ 1397 Hz). Since the experiment was designed as a multifeature MMN paradigm, several types of deviants were inserted in the melodies. Relevant for this study are pitch (or mistuning) deviants, which consisted of a 50-cents (quarter-tone) pitch rise with respect to the standard tones. See Quiroga-Martinez et al. (2019b) for further details.

### 2.3. A computational model of auditory expectation

To obtain continuous measures of auditory surprise we used IDyOM (Pearce, 2005, 2018), a variable-order Markov model of expectation that quantifies surprise as information content (IC):

Here *p* is the conditional probability of the current sound event, given previous events in the sequence and the long-term training of the model. This formulation implies that the lower the probability of the event, the larger its IC value and thus the surprise it generates in the listener. IDyOM estimates continuation probabilities by keeping track of the number of times a given pattern of events has occurred. The model takes into account patterns of variable length (“*n*-grams”) whose probabilities are combined through a smoothing process to produce the output values (see Pearce, 2005 for details). An advantage of IDyOM is that it can simulate different types of expectations. Specifically, it has a short-term (*stm*) submodel that generates dynamic expectations derived from the current auditory sequence and a long-term (*ltm*) submodel that simulates life-long schematic expectations derived from a large training corpus. In Omigie et al., (2013), a configuration that combined both the *ltm* and *stm* submodels was used. This configuration, known as *both+*, simulates a listener who employs short-term and long-term expectations to predict pitch continuations, but who also updates the *ltm* submodel with the knowledge gathered from the current stimulus (as indicated by the “+” symbol). Here, the corpus used to train the *ltm* submodel was the same as in Omigie et al. (2013) and corresponded to a collection of hymns and folk songs belonging to the Western tonal tradition.

IDyOM can use representations of different features of the auditory input, known as *viewpoints*, to make its probabilistic predictions. Following Omigie et al. (2013) and other behavioral work (Agres, Abdallah, & Pearce, 2018; Hansen & Pearce, 2014; Hansen et al., 2016), we used a joint representation of scale degree (“*cpint*”) and pitch interval (“*cpintfref*”) viewpoints to predict pitch (“*cpitch*”) continuations in the melodies. This is our reference model. The scale degree viewpoint assigns a category to each tone in a musical scale with reference to its tonal center, irrespective of its absolute pitch height. Scale degrees are hierarchically organized in Western tonal music so that, if context is kept constant, more prominent degrees (e.g., the tonic) are more frequent and are perceived as more expected than less prominent degrees (e.g., the leading tone) (Krumhansl, 1990). Transitions between scale degrees are idiomatic so that, for example, a leading tone would most often be followed by the tonic (Huron, 2006, p. 160). The pitch-interval viewpoint, on the other hand, quantifies the distance in semitones between consecutive tones. Idiomatic preferences pertaining to pitch interval also exist in Western tonal music such as the preponderance of small pitch intervals and note repetitions (Huron, 2006, p. 74).

To assess the contribution of different components to model performance, we compared different configurations of IDyOM. As a first step, we compared the reference configuration (*both+* with scale degree and pitch interval viewpoints combined) with a model that did not derive long-term knowledge from the current sequence (i.e., *both*). This aimed to reveal the extent to which listeners updated their long-term knowledge based on the current melodies. In a second step, the parameters were manipulated along two dimensions. First, we created submodels with either an *stm* or *ltm* component only, the comparison of which indicated the extent to which listeners relied on long-term or short-term expectations only. Second, we created submodels with either a scale degree or a pitch interval viewpoint only. Comparing these models allowed us to assess the extent to which participants’ expectations were primarily based on one of these musical features compared to the other. Orthogonal manipulations of these dimensions gave rise to the configurations shown in Figure 4 and table 3. IC values based on each of these configurations were obtained for every tone in the melodies. The values were binned into ten quantiles corresponding to increasing IC levels across the entire stimulus set and used for the analyses of the MEG data. Note that an *stm* model with scale degree and interval viewpoints was not included, as the range of IC values was not sufficiently large to avoid having the same value repeated in different quantiles.

**Table 2.** Widely Applicable Information Criteria (WAIC) and their standard error (SE), for each of the estimated mixed-effects models. For each comparison between adjacent models (i.e. “Model” vs “Null”), Bayes factors in favor of the alternative (BF_10_) and the null (BF_01_) are also reported. The predictors included in each model are indicated as follows: Int = Intercept, quant = IC quantile, hem = hemisphere, exp = expertise. Interactions are marked with colons (“:”). Note that the models also included participant-wise random effects for the intercept, the effects of IC quantile and hemisphere, and the quantile-by-hemisphere interaction. Comparisons with moderate or strong evidence for either the null or the alternative are highlighted in bold and marked with an asterisk “*”.

| Model | Predictors | Null | WAIC | WAIC (SE) | $BF_{10}$ | $BF_{01}$ |
| --- | --- | --- | --- | --- | --- | --- |
| *m0* | Int | NA | 8945.24 | 56.57 | NA | NA |
| ***m1*** | **Int + quant** | ***m0*** | **8839.13** | **55.83** | **> 1000\*** | **< .001** |
| ***m2*** | **Int + quant + hem** | ***m1*** | **8360.36** | **65.14** | **> 1000\*** | **< .001** |
| *m3* | Int + quant + hem + exp | *m2* | 8360.44 | 65.12 | 0.34 | 2.93 |
| ***m4*** | **Int + quant + hem + exp + quant:exp** | ***m3*** | **8361.24** | **65.06** | **0.24** | **4.21\*** |
| ***m5*** | **Int + quant + hem + exp + quant:exp + quant:hem** | ***m4*** | **8362.08** | **64.69** | **<.001** | **434.07\*** |
| *m6* | Int + quant + hem + exp + quant:exp + quant:hem + hem:exp | *m5* | 8362.46 | 64.71 | 0.89 | 1.12 |
| *m7* | Int + quant + hem + exp + quant:exp + quant:hem + hem:exp + quant:hem:exp | *m6* | 8363.41 | 64.67 | 0.43 | 2.33 |

**Table 3.**
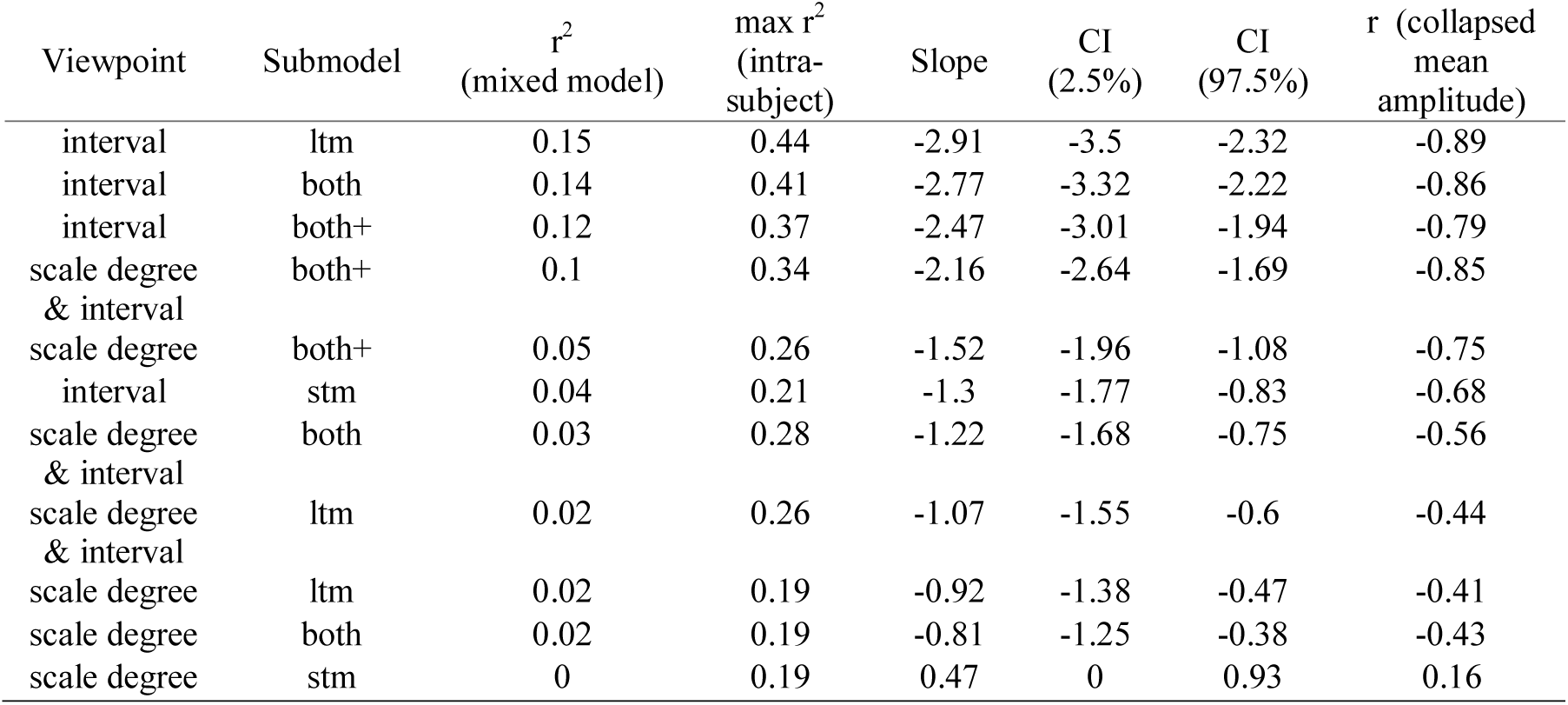
Performance of different IDyOM configurations in predicting the N1m amplitude from estimates of information content (IC), as measured by the variance explained at the group (mixed-model) and individual (intra-subject) level. The slope of the association between surprise (IC quantile) and N1m amplitude and corresponding confidence interval (CI), and the correlation between estimated surprise and the N1m amplitude averaged (or collapsed) across all subjects are also reported. The latter provides measures comparable to the correlations reported in Omigie et al. (2013).

**Figure 1.**
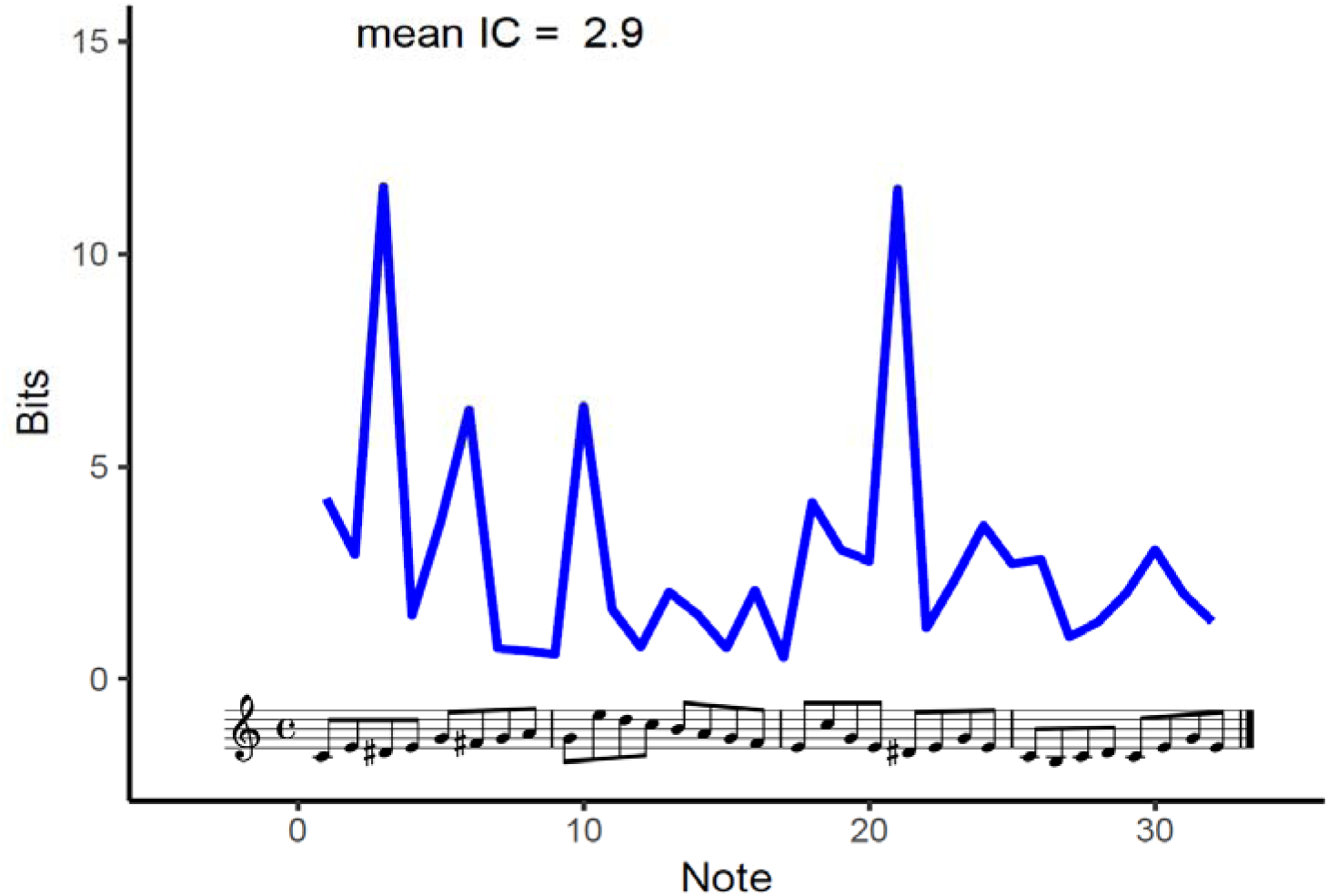
Example of one of the melodies used in the experiment and its estimated information content (IC, blue line). Values were obtained employing the reference model (*Both+* with scale degree and pitch-interval viewpoints combined). Note that out-of-key accidental tones (preceded by sharps, i.e. #) that belong to the 12-tone chromatic set (corresponding to all keys on the piano) but fall outside the 7-tone diatonic set (corresponding to the white keys on the piano in the key of C-major) typically result in higher information content (IC) than notes belonging to the diatonic set. For this experiment, out-of-tune deviant tones outside the chromatic set were also introduced (not displayed here). See Quiroga-Martinez et al. (2019b) for details in this regard, and the high-entropy (HE) condition in the supporting file 1 of the same study for the full stimulus set and corresponding IC estimates.

**Figure 2.**
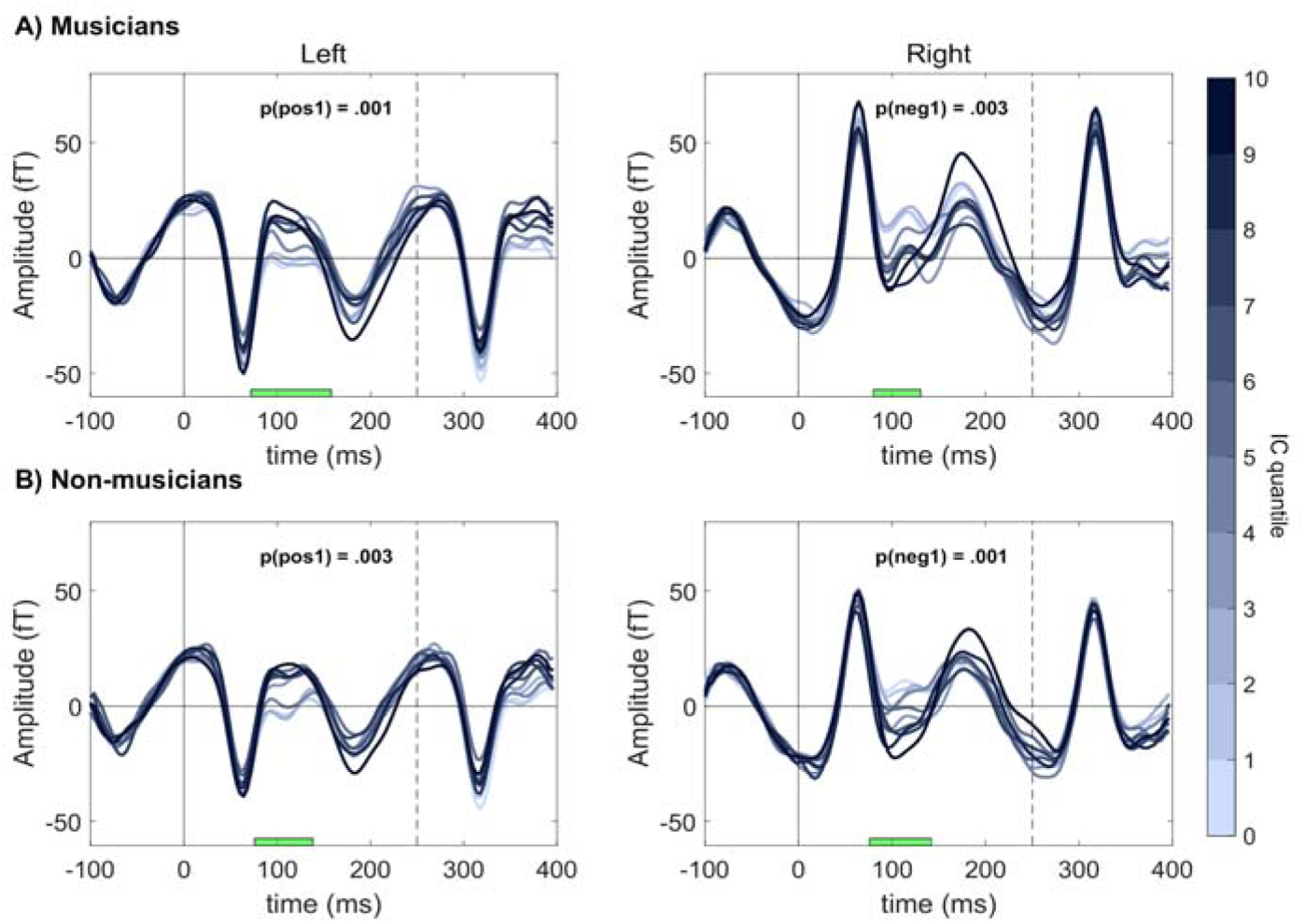
Effect of surprise (IC quantile) on the amplitude of the evoked response for both groups and hemispheres. For descriptive purposes, the time points when positive (“pos”) and negative (“neg”) clusters were significant are indicated with green horizontal bars. Note that this is not an accurate estimate of the true temporal extent of the effects (Sassenhagen & Draschkow, 2019). Displayed activity corresponds to the average of the four temporal magnetometers in each hemisphere with the strongest effect (left channels: 0231, 0241, 1611, 1621; right channels: 1341, 1331, 2411, 2421). Vertical dashed lines indicate the onset of the next tone.

**Figure 3.**
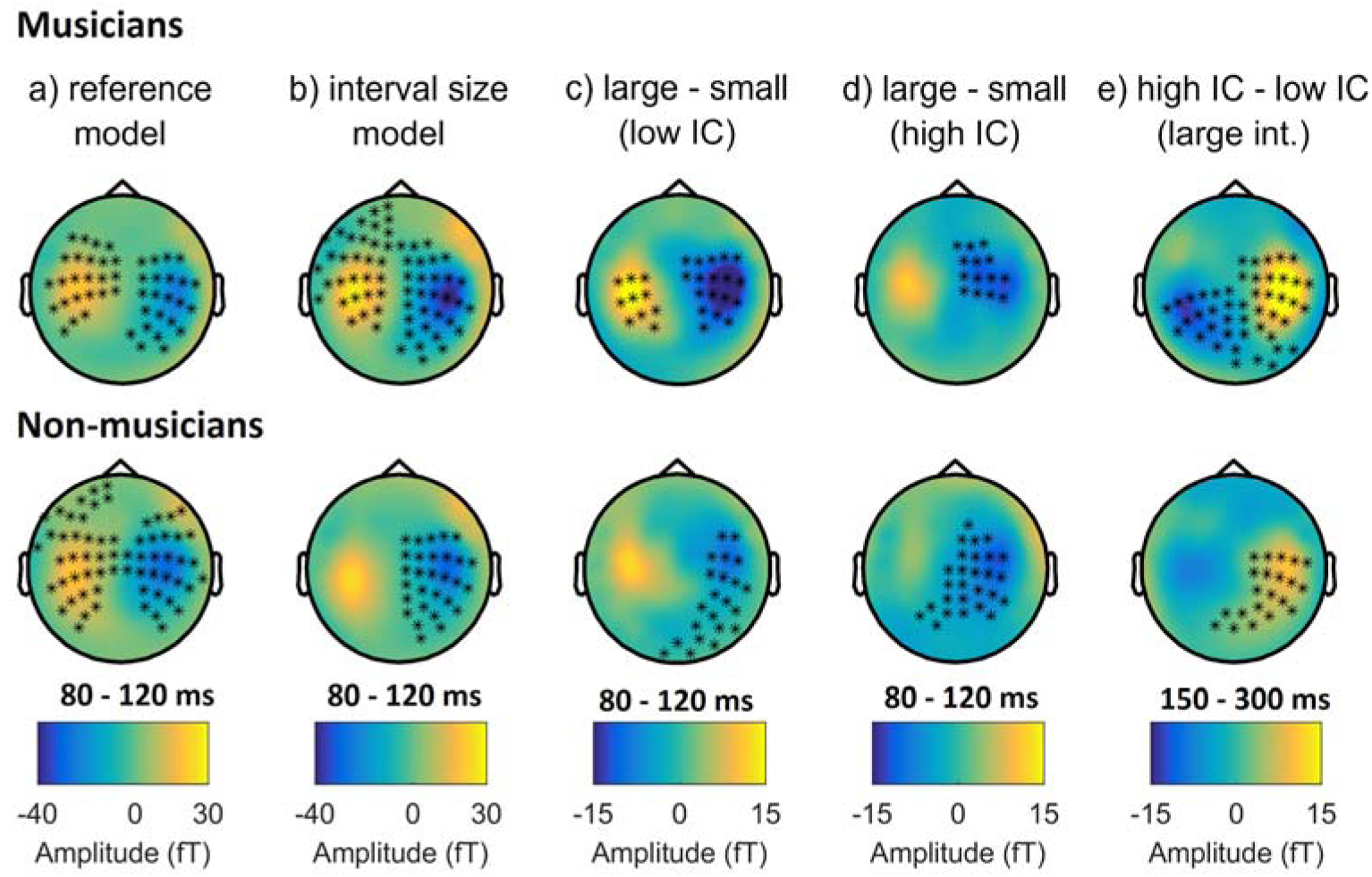
Topographic maps of the difference between: **a)** surprising (quantile 10) and unsurprising (quantile 1) tones for the reference model (*both*+ with scale degree and pitch interval viewpoints combined); **b)** tones following large (≥ 5 semitones) and small (1 semitone) intervals; tones following large (≥ 3 semitones) and small (≤ 2 semitones) intervals with either **c)** low-IC or **d)** high-IC; and **e)** high-IC and low-IC tones following large intervals. For **a, b, c**, and **d**, activity between 80 and 120 ms is displayed, corresponding to a modulation of the N1m component; whereas for **e**, activity between 150 and 300 ms is displayed, potentially corresponding to a P3am component. Note the change in polarity between the two time windows. Stars mark the channels where regression analyses (**a, b**) or pairwise contrasts (**c, d, e**) were significant. Note that this is not an accurate estimate of the true spatial extent of the effects (Sassenhagen & Draschkow, 2019).

**Figure 4.**
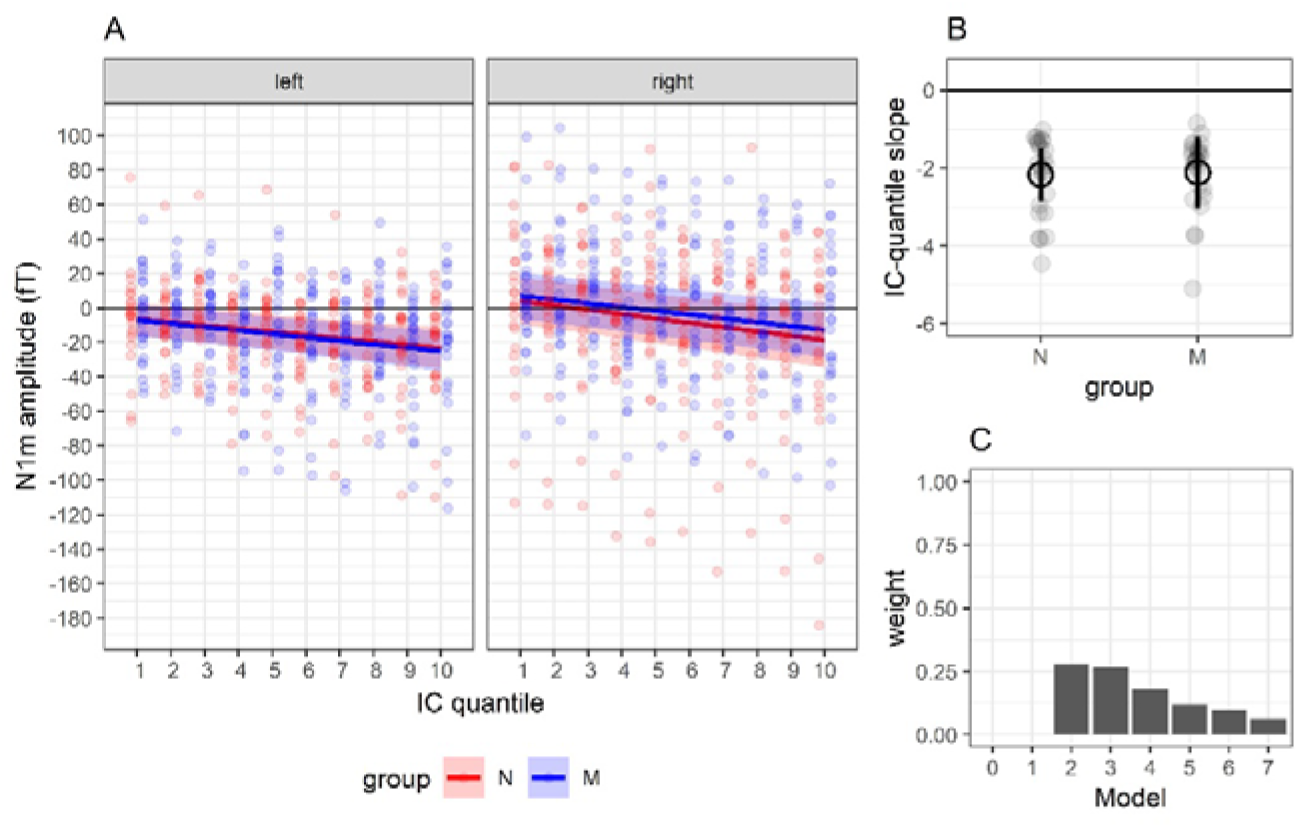
A) N1m amplitude as a function of surprise (IC quantile). Bayesian estimates of the IC quantile slopes (colored lines) and corresponding 95% credible intervals (shaded areas) are also displayed. Note that mean amplitudes in the left hemisphere were multiplied by −1 in order for them to have the same polarity as those in the right hemisphere. B) Bayesian estimates of the individual-(small gray dots) and group-level (larger circles) association between IC quantile and N1m amplitude. Error bars represent 95% credible intervals. C) Posterior model weights resulting from the incremental Bayesian model comparisons. N = non-musicians, M = musicians. See Table 2 for a description of each model.

### 2.4. Procedure

At the beginning of the session, participants received oral and written information and gave their consent. Then they filled out the Gold-MSI questionnaire and completed the MET. Once participants had put on MEG-compatible clothing, electrodes and coils were attached to their skin and their heads were digitized. During the MEG recording, they were sitting upright in the MEG device looking at a screen. Before presenting the musical stimuli, their auditory threshold was measured through a staircase procedure and the sound level was set at 60 dB above threshold. Participants were instructed to watch a silent movie, ignore the sounds, and move as little as possible. They were told there would be music playing in the background interrupted by short pauses so that they could take a break and adjust their posture. Sounds were presented through isolated MEG-compatible ear tubes (Etymotic ER•30). The MEG recording lasted approximately 90 minutes and the whole experimental session took between 2.5 and 3 hours including consent, musical expertise tests, preparation, instructions, breaks, and debriefing.

### 2.5. MEG recording and preprocessing

Magnetic correlates of brain activity were recorded at a sampling rate of 1000 Hz using an Elekta Neuromag MEG TRIUX system with 306 channels (204 planar gradiometers and 102 magnetometers). Participants’ head position was continuously monitored with four coils (cHPI) attached to the forehead and the mastoids. Offline, the signals coming from inside the skull were isolated with the temporal extension of the signal source separation (tSSS) technique (Taulu & Simola, 2006) using Elekta’s MaxFilter software (Version 2.2.15). This procedure included movement compensation in all but two non-musicians, for whom continuous head position information was not reliable due to suboptimal placement of the coils. However, in these cases the presence of reliable auditory event-related fields (ERFs) was successfully verified by visually inspecting the amplitude and polarity of the P50m component. Eye-blink and heartbeat artifacts were corrected with the aid of electrocardiography (ECG) and electrooculography (EOG) recordings, and independent component analysis as implemented by a semi-automatic routine (FastICA algorithm and functions *find_bads_eog* and *find_bads_ecg* in the software MNE-Python) (Gramfort, 2013). Visual inspection was used as a quality check.

The ensuing analysis steps were conducted with the Fieldtrip toolbox (version r9093) in Matlab (Oostenveld, Fries, Maris, & Schoffelen, 2011). Epochs comprising a time window of 0-400 ms after sound onset were extracted and baseline-corrected, with a pre-stimulus baseline of 100 ms. The epochs were then low-pass filtered with a cut-off frequency of 35 Hz and down-sampled to a resolution of 256 Hz. Each epoch was assigned to a category according to the IC quantile of the corresponding sound, for a given IDyOM model. For each participant and quantile, ERFs were computed by averaging the responses for all the tones belonging to the quantile. The four initial tones of each melody were excluded from the analyses to avoid neural activity related to the transition between melodies (e.g., effects of key change). Tones preceded by a deviant were also excluded to avoid carryover effects from the deviant response. Between 250 and 350 epochs were averaged per IC quantile.

### 2.6. Sensor-level analyses

The statistical analyses were performed on the magnetometers because this allowed us to properly look at the polarity of the magnetic fields, which is fundamental to disentangle different components such as the P50m, N1m and P2m. To assess whether ERF amplitude increased with information content, we performed a dependent-samples regression (*ft_statfun_depsamplesregrT* function in Fieldtrip) for musicians and non-musicians separately in a mass-univariate analysis. This type of regression employs the mean of the participant-wise coefficients to assess whether the association is greater than zero, thus giving a group-level *t*-statistic as output. To account for multiple comparisons, we used cluster-based permutations with *p* = 0.05 as cluster-forming threshold, t-*maxsum* as statistic, and 10,000 iterations. A time window between 0 and 300 ms after onset was selected, to avoid substantial overlap with the neural activity of the next tone (starting at 250 ms), but at the same time explore possible unexpected effects (e.g., in the P50m or the P2m components). To assess whether the association between neural activity and surprise was different for musicians and non-musicians, in a first-level analysis we obtained *t-*values reflecting the strength of association between IC quantile and neural activity for each participant, at each time point and sensor (*ft_statfun_indepsamplespregrT* function), which were then compared between groups in a second-level analysis with a two-sided independent-samples *t*-test (*ft_statfun_indepsamplesT* function). Cluster-based permutations were used to correct for multiple comparisons as indicated above.

To assess the relative evidence for the absence or the presence of an IC quantile-by-expertise interaction, we applied Bayesian estimation to mean N1m amplitudes. These amplitudes corresponded to the mean activity *±* 20 ms around the peak, extracted from the average of the 4 channels in each hemisphere that showed the strongest effect of IC quantile in the mass-univariate analyses (left channels: 0231, 0241, 1611, 1621; right channels: 1341, 1331, 2411, 2421). Since the ERFs have opposite polarities in different hemispheres, we multiplied left-hemisphere mean amplitudes by −1, so that they had the same (negative-going) polarity found in right-hemisphere amplitudes.

Several mixed-effects models were estimated using the *brm* function from the *brms* package (Bürkner, 2017) in R (R Core Team, 2019) and compared in an incremental way, adding one term at the time until reaching a full model with all main effects and interactions (Table 2). Participant random effects were included for intercept and slopes. The prior for the effect of IC quantile was taken from the reported results in Omigie et al. (2013), which showed a difference of around 1.5 μV between quantiles 1 and 10 for control participants. Standardizing this difference with the reported variance gives an effect size of about 0.96. This effect was extrapolated to our magnetometer data using the variance around the N1m time window taken from a previous auditory dataset collected with the same MEG scanner, which resulted in a difference of about 30 fT between quantiles 1 and 10. Therefore, a plausible slope would be 30/9 = 3.33 fT/quantile, which we rounded to 3.5 fT/quantile. With this rough estimate, we set a conservative Gaussian prior centered at 0 with SD = 3.5 fT/quantile. In other words, we regard effects similar or smaller than 3.5 fT/quantile as most likely, and effects larger than twice this value as very unlikely. For the IC quantile-by-expertise interaction we set a similar prior centered at zero with SD = 1.75 fT/quantile, which means that we regard modulations equal to or smaller than half of the main effect to be most likely. The same prior was set for other interaction terms involving IC quantile. Regarding the main effects of expertise and hemisphere, Gaussian priors centered at zero with SD = 10 fT were set, which are also conservative and represent rather small effect sizes. Finally, for the intercept of the model as well as the standard deviation and random effects, weakly informative priors were set, corresponding to a uniform distribution between 0 and 100 fT. Models were built in an incremental way, with *m0* having an intercept only (null model), *m1* adding a term for IC quantile, *m2* adding a term for hemisphere, and so on (Table 2). The full model (*m7*) included all main effects and interactions. Comparisons between adjacent models (i.e., models with and without a particular factor) were performed by estimating Bayes factors (BF) and Widely Applicable Information Criteria (WAIC) (Wagenmakers et al., 2018; Watanabe, 2010). Of particular interest is the comparison between *m3* and *m4*, as it assessed the evidence for the IC quantile-by-expertise interaction.

#### 2.6.1. Comparisons between IDyOM configurations

To compare the different IDyOM configurations shown in Table 3, several metrics were used. First, we took the peak *r*^*2*^-value across the whole sensor array for each model, resulting from the average of participant-wise first-level regression analyses (*ft_statsfun_indepsamplesregrT* function). Second, we obtained maximum likelihood estimates of linear mixed effects models (function *lmer*, package *lme4*, Bates, Mächler, Bolker, & Walker, 2015) of N1m mean amplitudes (instead of Bayesian estimates for speed of processing), including IC quantile and hemisphere as predictors, and random intercepts and slopes for participants. *r*^*2*^-values for the effect of IC quantile were also obtained.

#### 2.6.2. Comparison with the MMNm

The preprocessing steps for the measurement of the MMNm were the same as for the N1m, with the difference that, instead of surprise levels, an ERF was computed for only two conditions: standards and deviants. The MMNm is the difference between these two conditions, which was assessed here for magnetometers through paired-samples *t*-tests in mass-univariate analyses (note that statistical analyses for the MMNm had previously been conducted for gradiometers in Quiroga-Martinez et al. 2019a, 2019b). The deviants analyzed here correspond to out-of-tune tones (i.e., tones with pitches outside the musical tuning system that cannot normally be played on a piano). Note that other deviants were also present in the experiment, but we focused on mistunings as they are disruptive with regard to pitch, which is the feature modeled with IDyOM in this study. Independent-samples *t*-tests on MMNm difference waves were used to assess the effect of expertise. For the comparison between the two components, the N1m was calculated as the difference between IC quantile 10 and IC quantile 1. In this case, the model with the best performance was used (*ltm* with a pitch-interval viewpoint), to increase the signal-to-noise ratio. Two-sided paired-samples *t*-tests were used to compare the resulting MMNm and N1m difference waves. Cluster-based permutations were employed as multiple comparisons corrections in all analyses, as described above. Since, based on visual inspection, the peak latency of the N1m and the MMNm seemed to be different—which might give misleading results in the permutation tests—we additionally performed two-sided paired-samples *t*-tests on peak latencies and mean amplitudes separately, estimated as described above.

### 2.7. Source reconstruction

For source reconstruction we employed the multiple sparse priors method (Friston et al., 2008) in SPM12 (version 7478). Individual anatomical magnetic resonance images (MRI) were available for 20 musicians and 20 non-musicians only. These are the participants included in the analysis. In the case of two excluded musicians and one excluded non-musician, the images were corrupted by artefacts. The remaining excluded participants did not attend the MRI session. Two brain images were acquired with a magnetization□prepared two rapid gradient echo (MP2RAGE) sequence (Marques et al., 2010) in a Siemens Magnetom Skyra 3T scanner. These images were combined and motion-corrected to form unified brain volumes, which were subsequently segmented, projected into MNI coordinates, and automatically coregistered with the MEG sensor array employing digitized head shapes and preauricular and nasion landmarks. Coregistration outputs were visually inspected. Lead-fields were constructed using a single-shell BEM model with 20.484 dipoles (fine grid). A volume of the inverse solution was created for each participant between 75 and 125 ms for the N1m (difference between quantile 10 and quantile 1 for the *ltm* pitch-interval model), and between 175 and 225 ms for the MMNm (difference between deviants and standards). These time windows were chosen based on the peak amplitude of each component and are warranted by the output of statistical tests. The volumes for each component were submitted to a two-sided one-sample *t*-test to reveal the sources consistently identified across all participants. To assess possible differences between the N1m and the MMNm, the volumes were also compared in a two-sided paired-samples *t*-test. The error rate of voxel-wise multiple comparisons was corrected with random field theory, with a cluster-level alpha threshold of 0.05 (Worsley, 2007).

## 3. Results

### 3.1. Modulation of the N1m and expertise effects

Dependent-samples regressions revealed a significant association between the amplitude of the evoked responses and IC quantile (Figure 2) for both musicians and non-musicians around 100 ms after sound onset. This association was largest at temporal magnetometers, was negative in the right hemisphere and was positive in the left hemisphere (Figure 3a). This reflects the inversion of the recorded magnetic field given the hemisphere-dependent orientation of the source with respect to the sensor array. No significant differences between musicians and non-musicians were detected in the second-level analysis (smallest *p*-value = .2).

### 3.2. Bayesian analyses of N1m amplitudes

Results from the Bayesian analyses are shown in Table 2 and Figure 4. WAIC values indicate that adding IC quantile (*m1*) and hemisphere (*m2*) significantly improves predictive power, but adding the main effect of expertise (*m3*) or any interaction term does not improve the predictive power any further. Estimations of model weights (Figure 4c) also suggest *m2* as the winning model. A similar picture can be inferred from Bayes factors. The *m1* model**—**with a term for IC quantile**—**is much more likely than an intercept-only model (*m0*), and the *m2* model**—**including hemisphere**—**is much more likely than the *m1* model. This strong effect of hemisphere might be a carryover from the P50m component, which was larger in right-hemisphere channels. No other comparisons support alternative models. Instead, in a few cases there is evidence for null models. Of paramount interest here is the comparison between *m3* and *m4*, which suggests that a model with no interaction between IC quantile and expertise is about 4.2 times more likely than a model with it (Table 2). We regard this as moderate evidence for the null hypothesis. Similarly, a model with no IC quantile-by-hemisphere interaction (*m4*) is about 434 times more likely than a model with it (*m5*). The remaining BF suggest the absence of an effect of expertise, the absence of a hemisphere-by-expertise interaction, and the absence of a three-way-interaction between IC quantile, hemisphere and expertise. In these cases, however, the BF are inconclusive. Finally, a look at the parameters shows that only the estimates for the main effects of IC quantile and hemisphere are credibly different from 0 across all models compared (supplementary file 2).

### 3.3. Comparison of IDyOM configurations

The performance of IDyOM configurations in terms of predicting the N1m amplitude from IC estimates is reported in Table 3. Even though the reference model (*Both+*, with scale degree and interval viewpoints combined) performed well, models that included only a long-term component and an interval-only viewpoint performed better. This was reflected in larger *r*^2^-values and steeper slopes. Models with only scale-degree viewpoints and/or short-term components performed poorly. The best model employed a long-term component and an interval-only viewpoint. Given that in Western tonal music (and therefore in our training corpus), smaller intervals are more common than larger ones, it could be the case that the effects observed are caused by the size of the interval itself rather than its IC, which would explain why long-term interval-only models had the best fit. For this reason, in further exploratory analyses, four categories of tones were averaged for each participant, comprising small-interval transitions (≤ 2 semitones) with either low IC (small/low-IC, ≤ 4) or high IC (small/high-IC, > 4); and large-interval transitions (≥ 3 semitones) with either low IC (large/low-IC) or high IC (large/high-IC). Cluster-based, permutation-corrected paired-samples *t*-tests conducted in a 50-300 ms time window revealed significant differences between large/high-IC and large/low-IC for musicians and non-musicians (Figure 6). Crucially, these differences were found in later time windows that excluded earlier N1m latencies. Moreover, the direction of this effect was the opposite of that of the N1m modulation. This resulted in the polarity shown in Figure 3e, which would presumably correspond to a positive deflection in an EEG recording and could be interpreted as the magnetic counterpart of the P3a (P3am) (see section 4.2). Using the methods described above, the neural generators of this effect were localized in frontal (peak location: 42, 32, −14), parietal (peak locations: −36,−64, 54; 56, −24, 32 and 38,−26, 56) and inferior temporal regions (peak location: 52, - 58, −6) (Figure 8). Note, however, that the signal-to-noise ratio of this effect was lower than that of the MMNm or the N1m effect, which resulted in no significant differences after multiple-comparisons correction. For this reason, we report uncorrected statistical maps thresholded at p < 0.0005.

**Figure 5.**
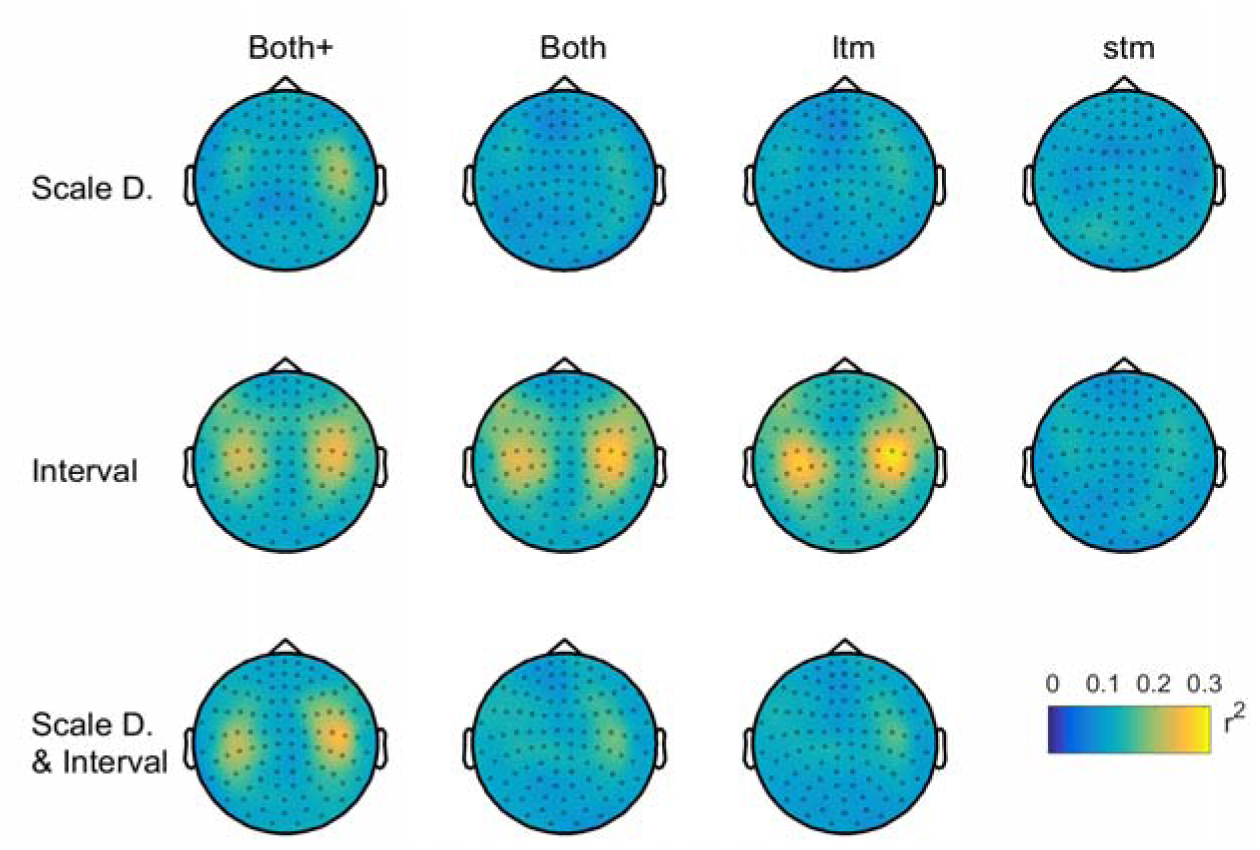
Grand-averaged topographic maps of the variance (*r*^2^) in neural activity explained by each IDyOM configuration for each participant (i.e., first-level intra-subject variance). Configurations included scale degree and pitch interval viewpoints, and their combination, as well as short-term (*stm*) and long-term (*ltm*) submodels, and their combination (*Both* and *Both+*).

**Figure 6.**
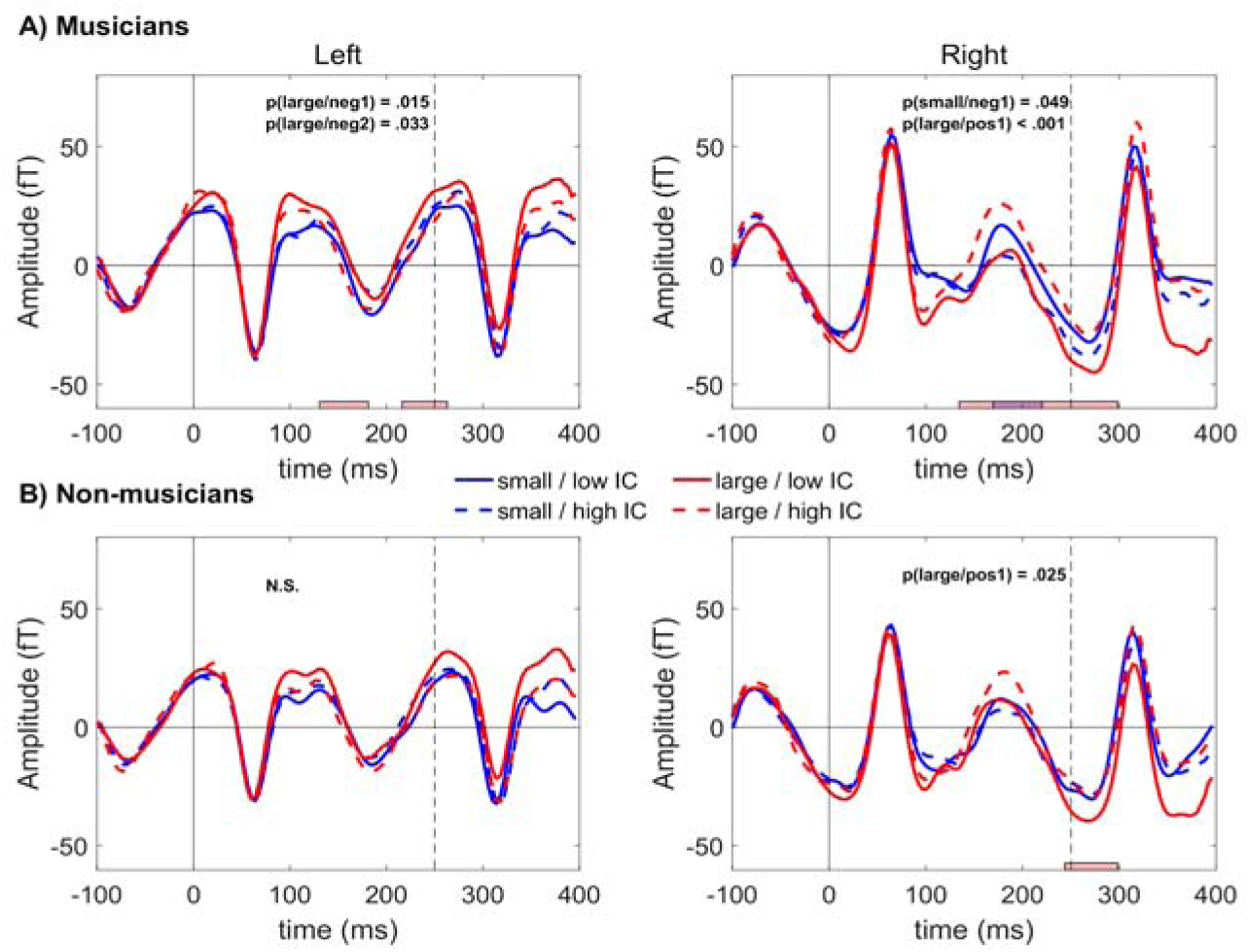
Event-related fields for tones with high (> 4 bits) or low (≤ 4 bits) surprise (i.e., information content) following small (≤ 2 semitones) or large (≥ 3 semitones) intervals. The reference model (*Both*+ with scale degree and pitch-interval viewpoints combined) was used to obtain the estimates. For descriptive purposes, horizontal colored lines indicate positive (“pos”) or negative (“neg”) clusters associated with differences between conditions. Note that they are not an accurate estimate of the true temporal extent of the effects (Sassenhagen & Draschkow, 2019). The color of the lines indicates whether the contrast between high-IC (dashed lines) and low-IC (solid lines) tones was made for large (red) or small (blue) intervals. Vertical dashed lines mark tone onsets.

**Figure 7.**
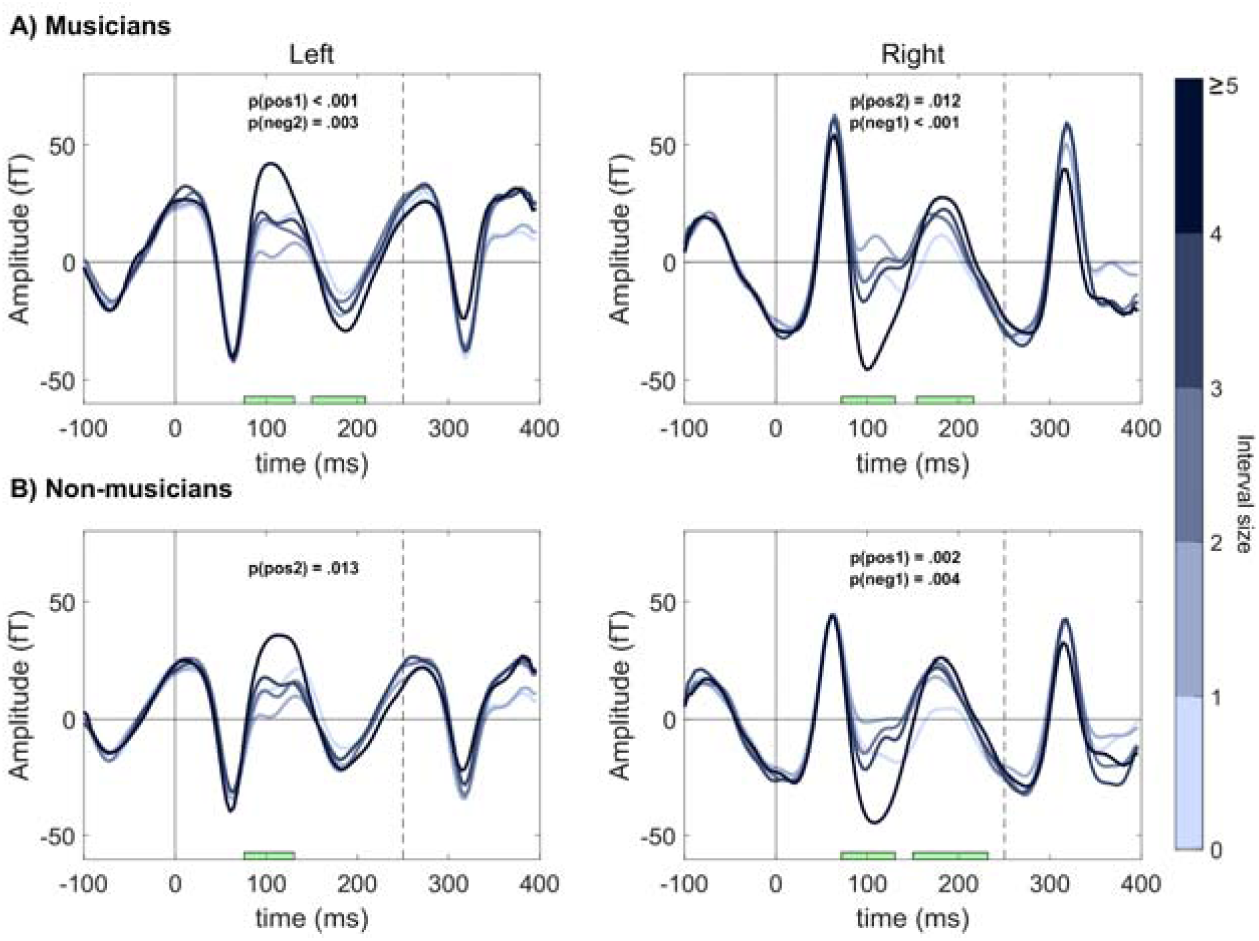
Neural responses to pitch intervals of different size (in semitones) in both groups and hemispheres. For descriptive purposes, the time points when positive (“pos”) and negative (“neg”) clusters were significant are indicated with green horizontal bars. Note that they are not an accurate estimate of the true temporal extent of the effects (Sassenhagen & Draschkow, 2019). Displayed activity corresponds to the average of the four temporal magnetometers in each hemisphere with the strongest effect (left channels: 0231, 0241, 1611, 1621; right channels: 1341, 1331, 2411, 2421). Vertical dashed lines indicate the onset of the next tone.

**Figure 8.**
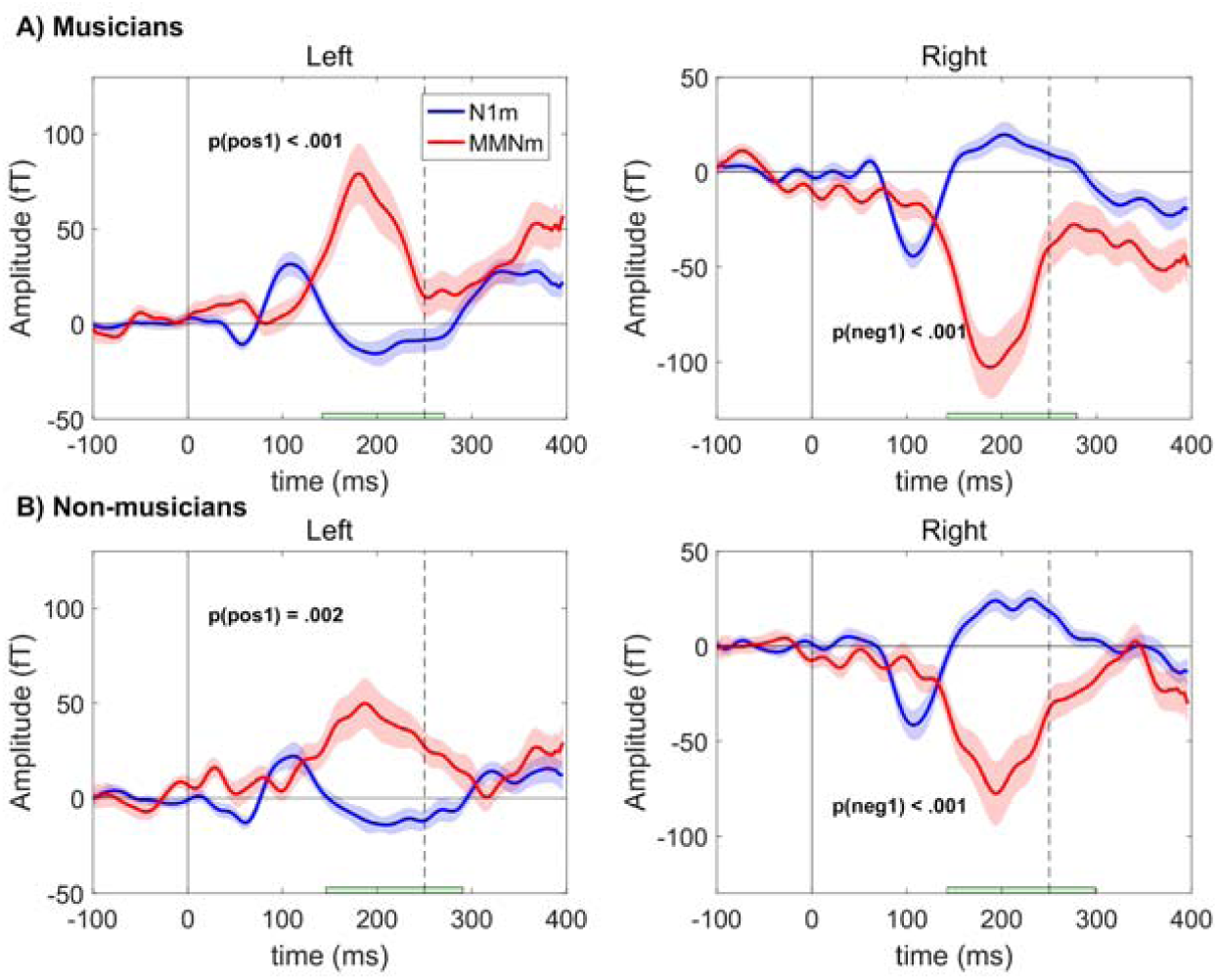
Event-related fields of the N1m effect (i.e. quantile 10 - quantile 1 of the *ltm* interval-only model) and the MMNm (deviant - standard). For descriptive purposes, green horizontal lines indicate the times when differences between components were significant. Note that this is not an accurate estimate of the true temporal extent of the effects (Sassenhagen & Draschkow, 2019). Shaded areas depict 95% confidence intervals. Displayed activity corresponds to the average of the four temporal magnetometers in each hemisphere with the strongest N1m modulation (left channels: 0231, 0241, 1611, 1621; right channels: 1341, 1331, 2411, 2421). Dashed vertical lines indicate tone onsets.

Regarding small intervals, a significant difference was found between small/high-IC and small/low-IC around 200 ms after sound onset, only for musicians in the right hemisphere. The direction of this effect was opposite to the one found for the comparison between large/high-IC and large/low-IC (Figure 6). Note, however, that the *p*-value was close to the significance threshold in this case. Finally, in contrast to these analyses, when large and small intervals were compared for low-IC and high-IC tones separately, significant differences in the N1m time window (50-150 ms) in the expected direction were found in both groups (Figures 3c and 3d).

In addition, we performed a within-subjects regression analysis using absolute interval size (with no IDyOM modeling) as predictor. We grouped and averaged the tones into five categories comprising intervals with either 1, 2, 3, 4 or ≥5 semitones. The number of categories could not be increased as there were few instances of large intervals. The analysis showed that interval size was strongly associated with the amplitude of the N1m in musicians and non-musicians (Figures 7 and 3b; maximum grand-averaged intra-subject *r*^*2*^ = 0.52). The variance explained by this model was higher than the one explained by any of the IDyOM configurations (Table 3). However, note that part of this outcome might be due to including five instead of ten categories, which already reduced the variance in the data. Interestingly, all categories showed a monotonic amplitude increase, except for 1-semitone intervals, whose amplitude was larger than expected (Figure 7).

### 3.4. Comparison between the MMNm and the N1m difference wave

Mass-univariate analyses revealed that the MMNm and the N1m difference wave (high IC – low IC) were substantially different (Figure 8). The latency of the MMNm was significantly longer (*t* = 35.68, p < .001) and its amplitude was significantly larger (*t* = 8.37, p < .001) than those of the N1m effect, in the mean and peak amplitude analyses. Moreover, expertise effects were found for the MMNm (p < .001), thus reproducing in the magnetometer space the differences previously reported for combined gradiometers (Quiroga-Martinez et al. 2019a). Regarding source analyses, one-sample *t*-tests suggested that the main generators of both the MMNm and the N1m effect were located in the surroundings of primary auditory cortex, as expected (Figure 9). The peak activation for the MMNm was located in the anterior part of the primary auditory cortex (A1) in the right hemisphere (56, 0, 2), and in the posterior portion of A1 in the left hemisphere (−48,−16, −4). For the N1m effect, the peak was located in the posterior part of A1 in both hemispheres (right: 48, −18, 6; left: −46,−18, −4). The contrast between the two components did not yield significant results.

**Figure 9.**
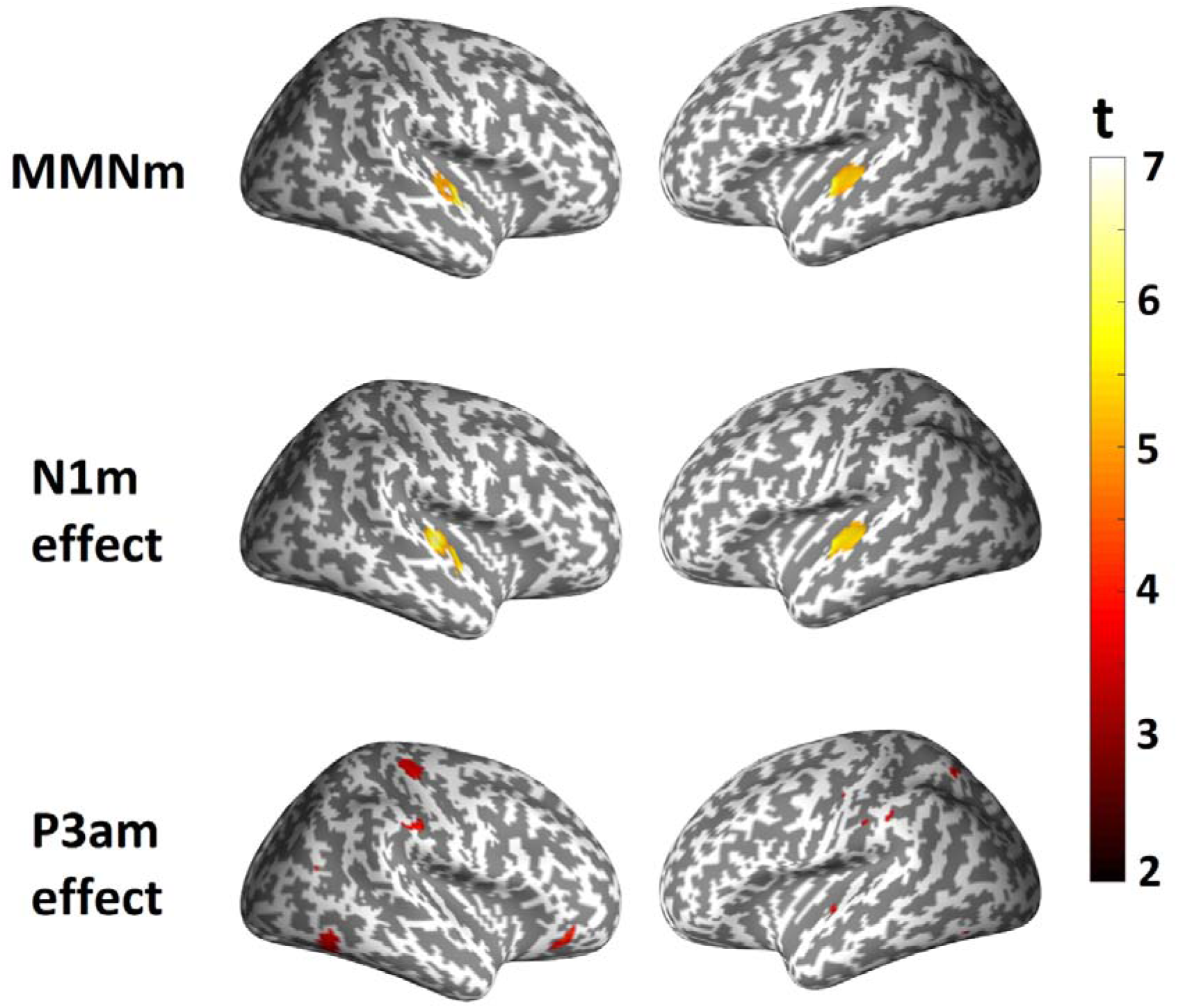
Neural generators of the MMNm, N1m effect and P3am effect. Color maps depict *t*-statistics from one-sample *t*-tests. For the N1m and MMNm, the maps are thresholded at *p* = 0.05, after multiple-comparisons correction. For the P3am effect the map is thresholded at p = 0.0005, without multiple-comparisons correction. Participants from both groups were included in the tests, as no statistically significant differences were detected between musicians and non-musicians.

## 4. Discussion

In this study, we aimed to determine whether the modulation of neural activity by subtle changes in auditory surprise—as estimated with a computational model of expectations (IDyOM)—is affected by musical expertise. Our results showed that surprise estimates were associated with the amplitude of the N1m in both groups, thus replicating the findings by Omigie and colleagues (2013). However, contrary to our expectations, non-significant null-hypothesis-testing results and Bayesian analyses indicated no differences between musicians and non-musicians in the strength of this association.

### 4.1. Interval size and sensory adaptation better explain N1m responses

Further exploratory analyses provided clues about the nature of the expectations reflected in the N1m modulation, and an explanation for the lack of differences between the groups. Comparisons of IDyOM configurations indicated that models predicting pitch interval based on long-term knowledge performed best, and even better than the reference model used by Omigie and colleagues (2013) (*Both+* with pitch interval and scale degree viewpoints combined). In contrast, models predicting scale degree based on short-term expectations performed the worst. Given that, in Western tonal music (which is the style that IDyOM was configured to model in the present experiment), smaller intervals are much more common and therefore overall less surprising than larger intervals, it is possible that the modulation of the N1m could be explained by a much simpler factor: interval size.

In additional analyses, we showed that if high-IC (i.e., surprising) tones are contrasted with low-IC (i.e., less surprising) tones when interval size is kept constant, the differences previously observed in the N1m are not detected anymore. Conversely, if tones following a large interval are contrasted with those following a small interval while surprise is kept constant, then the differences still persist. Furthermore, N1m amplitude was shown to significantly increase with larger interval sizes. All of this indicates that interval size, rather than probabilistic prediction, is the most likely factor behind the reported effect.

This explanation is compelling when one considers that the amplitude of the N1 is modulated by stimulus specific adaptation or SSA (May & Tiitinen, 2010; May, Westö, & Tiitinen, 2015; Näätänen & Picton, 1987; Pérez-González & Malmierca, 2014). SSA occurs when neural populations that respond to the spectral content of the stimulus become less responsive with subsequent presentations of the same or a similar stimulus, likely due to synaptic depression (May et al., 2015; Ulanovsky, 2004; Ulanovsky, Las, & Nelken, 2003; Yarden & Nelken, 2017). Thus, tonotopic neurons responding to a tone become adapted, and therefore the neural activity generated by an equal or spectrally similar successive tone would be attenuated. However, if the successive tone is sufficiently different, non-adapted neurons would be engaged, thus producing more robust neural responses. Therefore, while the melodies used here were deliberately composed to avoid pitch repetitions, it could be that our results are driven by increased neural responses to sounds farther apart in their spectral content, which is the case for tone transitions with larger pitch intervals. It has to be noted that, although SSA has almost exclusively been studied with pure tones—as opposed to the complex tones used here—some studies have demonstrated SSA for complex sounds (Nelken, Yaron, Polterovich, & Hershenhoren, 2013) and even frequency selectivity for broadband sounds (Rauschecker, Tian, & Hauser, 1995). Moreover, an acoustic analysis of our stimulus set (supplementary file 3) shows that the spectral similarity within pairs of piano tones decreases with pitch distance, which is consistent with the suggested explanation. Finally, pitch distance might not be the only relevant variable affecting SSA, as 1-semitone intervals were shown to have a larger amplitude than expected. This points to future research efforts in which the spectral difference between consecutive tones and its effect on N1 amplitudes are carefully modeled, measured and tested.

The results in Omigie et al. (2013) are in agreement with the explanation above. When comparing different IDyOM configurations, they found that those including a pitch-interval viewpoint performed the best. Note, though, that scale degree viewpoints had a good performance as well. This could be explained by the fact that scale degree and interval size are also correlated, since scale degree transitions are dominated by stepwise motion (Huron, 2006, p. 160). In our study, scale degree was also associated with neural activity, although to a lesser degree than in Omigie et al. (2013). This might have to do with potential differences in interval distributions between the two stimulus sets. Moreover, Hansen and Pearce (2014) found that long-term models were the best at predicting expectedness ratings. This is consistent with the *ltm*/pitch-interval configuration yielding the best performance in our analyses. This might be due to a more accurate estimation of pitch intervals in a Western tonal context—where interval size and interval probability are confounded—presumably arising from the larger amount of data present in the long-term training corpus compared to the stimuli themselves.

The current explanation is also consistent with early research showing that the amplitude of the N1 is larger when a pure tone is farther apart from preceding pure tones in the frequency continuum (Butler, 1968; Näätänen et al., 1988; Picton, Woods, & Proulx, 1978). Thus, our results generalize this effect to the spectral content of complex tones in more realistic and complex auditory sequences. Moreover, speech research showing modulations of the N1m component by the acoustic properties of phonemes provides further support to our proposal (Manca & Grimaldi, 2016; Shestakova, Brattico, Soloviev, Klucharev, & Huotilainen, 2004). On the same line, in a previous EEG study reporting an effect similar to the one found here, manipulations of surprise were also confounded with interval size, which is consistent with our data (Koelsch & Jentschke, 2010). Finally and most importantly, the proposed confound would explain why no differences between the groups were found in our study, since the long-term expectations captured by IDyOM rely on culture-specific knowledge of higher-order statistical dependencies between tones, which are arguably different from the early and rather unspecific sensory processes reflected in the N1m.

### 4.2. Hierarchical auditory predictive processing

The current results are not necessarily at odds with IDyOM being a good model of auditory expectation, for which there is otherwise solid supporting behavioral evidence (Agres et al., 2018; Bianco et al., 2019; Hansen & Pearce, 2014; Hansen et al., 2016; Morgan, Fogel, Nair, & Patel, 2019; Pearce, 2018). Rather, they indicate that probabilistic prediction is not reflected in the N1, but instead might engage later processing stages. Interestingly, in the analyses keeping interval size constant, IC modulated the amplitude of a later component in the case of large intervals. Given its latency, polarity and putative neural generators, this effect could be interpreted as a P3, a component associated with the orientation of attention and the engagement of higher-order cognitive processes (Masson & Bidet-Caulet, 2019; Polich & Criado, 2006; Squires, Squires, & Hillyard, 1975). Notably, in the first EEG study using IDyOM as a model of expectedness, surprising tones generated a similar positive late response that was larger for unexpected than expected sounds (Pearce, Ruiz, Kapasi, Wiggins, & Bhattacharya, 2010).

Modulations of the P3 have been found in global-local MMN paradigms where local expectations about tones and global expectations about patterns are generated and violated independently (Bekinschtein et al., 2009; Wacongne et al., 2011). In this case, P3 responses are observed for global deviants. Thus, it might be that the higher-order musical expectations reflected in IDyOM’s estimates are of a similar kind to the global patterns in the aforementioned paradigm. Note, however, that in Bekinschtein et al. (2009) responses to global deviants were observed only under conscious perception, which differs from the passive listening condition used in our study. Therefore, the late response found here may correspond to the magnetic counterpart of an early subcomponent of the P3, the P3a (or P3am), which is thought to reflect the orientation of attention to violations of the context, in contrast to the later P3b subcomponent, which is taken to reflect proper conscious perception (Polich & Criado, 2006; Squires et al., 1975).

In addition to the late effect of IC in the case of large intervals, we found similar differences for small intervals in musicians in the right hemisphere. However, the direction of this effect was opposite to that of the putative P3am, the reason for which is difficult to infer from our data. It might be the case that this effect corresponds to an emergent MMNm hidden by attention-related P3am responses in the case of large intervals, which are arguably more salient than small intervals. In any case, this effect should be interpreted with caution since the corresponding *p*-value was close to the significance threshold.

While these findings come from exploratory analyses and need to be replicated in a controlled experiment, they speak to a predictive processing hierarchy encompassing different representational levels and time scales. Thus, the N1m would be restricted to tonotopic predictions about the spectral content of sounds mainly driven by the immediate past. In contrast, later components, such as the P3am, may reflect higher-order stylistic predictions about categories derived from the initial sensory parsing—e.g., pitch—and would index the encoding of patterns spanning longer temporal scales. This is consistent with empirical evidence suggesting hierarchical processing along the auditory system (Escera & Malmierca, 2014; Griffiths & Warren, 2002; Parras et al., 2017; Rauschecker et al., 1995; Wacongne et al., 2011), and with theories of predictive processing in the brain (Clark, 2016; Friston, 2005; Friston, Rosch, et al., 2017; Vuust & Witek, 2014).

### 4.3. The distinction between MMN and N1 revisited

The relationship between N1 and MMN has been long debated (Jaaskelainen et al., 2004; May & Tiitinen, 2010; Näätänen, Jacobsen, & Winkler, 2005; Näätänen & Picton, 1987). While their scalp topographies and latencies are similar, and both components respond to changes in the auditory signal, they can be dissociated in specific cases. For example, an unexpected tone repetition (Tervaniemi, Saarinen, Paavilainen, Danilova, & Näätänen, 1994), tone omission (Bendixen et al., 2012), or intensity decrease (Näätänen et al., 2007) would attenuate the N1 but nonetheless elicit a robust MMN. This has led to the conclusion that the N1 is mainly modulated by SSA, whereas the MMN reflects the breach of a memory trace (Näätänen et al., 2007; Näätänen & Picton, 1987) or a probabilistic predictive model (Bendixen et al., 2012; Garrido et al., 2009; Lieder et al., 2013; Wacongne, Changeux, & Dehaene, 2012). This is consistent with the two components reflecting different levels of hierarchical processing.

Here, we were able to dissociate the N1m and the MMNm in the same subjects and in the same stimulus sequence. Consistent with hierarchical processing, our results indicate that the N1m effect is much smaller and happens earlier than the MMNm. Importantly, the sounds that gave rise to the MMNm in our experiment (i.e., quarter-tone mistunings) violated the musical tuning system, which prescribes the pitches and pitch intervals that can be expected (Brattico, Tervaniemi, Näätänen, & Peretz, 2006). This entails that the listener needs to infer abstract regularities and relationships between tones (such as: “the distance between consecutive pitches cannot be smaller than a semitone”), which goes beyond the sensory parsing reflected in the N1m. This is consistent with the fact that the MMNm, but not the N1m, was modulated by expertise, given that learning with precision the pitch heights and pitch intervals allowed in a musical system (i.e., learning to be “in tune”) is an essential musical skill.

Source reconstructions revealed overlapping generators for the N1m and MMNm in primary auditory cortex. This indicates that the two hierarchical processing stages are performed by the same or contiguous neural populations. Note that it has been previously argued that the neural generators of the MMN are slightly anterior to those of the N1 (e.g., Rosburg, Haueisen, & Kreitschmann-Andermahr, 2004; Sams, Kaukoranta, Hämäläinen, & Näätanen, 1991). These studies, however, typically rely on equivalent current dipoles for source estimation, whereas here we used distributed sources. Note, though, that peak activity in the right hemisphere was more anterior for the MMNm than the N1m, which would be consistent with the literature and with hierarchical processing stages. Methods such as intracranial recordings (e.g. Omigie et al., 2019) and dynamic causal modeling (Moran, Pinotsis, & Friston, 2013) could potentially disentangle the generators and the dynamics of the processes reflected in the two components, in the case of musical stimuli.

When considered together, our results present a coherent picture in which three stages of processing can be identified. In an initial stage, tonotopic neurons in A1 adapt to incoming sensory input thus modulating N1m amplitude. In a second stage, auditory objects are formed and low-level regularities are established, giving rise to the MMNm when violated. Finally, in a third stage, higher-order predictions arising from knowledge of the musical context are deployed. These are indexed by late components such as the P3am.

### 4.4. Implications, limitations and future directions

One limitation of our study is that the explanatory power of surprise estimates could not be assessed at the single-trial level due to the inherent noisiness of neurophysiological data. Nonetheless, this is an interesting future research direction that could benefit from techniques such as intracranial recordings and multivariate pattern analysis (e.g. Demarchi, Sanchez, & Weisz, 2019). Moreover, while here we found compelling evidence for the dissociation between interval size and estimated surprise, a more controlled study is needed to properly demonstrate this claim and confirm SSA as the underlying mechanism. Similarly, although we employed more realistic stimuli than many previous studies on auditory predictive processing, the melodies used here are still far from being truly musical. Future experiments could address, for example, how introducing rhythm, expressive timing and dynamics and concurrent melodic lines affects predictive processing at the sensory levels reflected in the N1. Another caveat is that information about musicians’ practiced musical style was not available. This variable could be relevant to rule out that the lack of differences between groups is simply due to the lack of familiarity of musicians with Western tonal music. However, this explanation is unlikely considering the widespread presence of Western tonal music; the fact that the experiment was conducted in a Western country; the typical repertoire associated with the instruments played by the musicians (supplementary file 1), and the strikingly similar association between IC and N1m amplitudes in both groups (Figure 4).

Despite these shortcomings, our findings have clear consequences for research on music and auditory predictive processing. We have shown that it is crucial to account for interval size and SSA when addressing statistical learning, even in the case of complex sounds. Regarding music perception, our results provide neural evidence that early sensory mechanisms might be fundamental for the perceptual processing of melodies and coexist with higher-order probabilistic predictions arising from knowledge of musical styles. Similarly, the evidence presented makes a contribution to the ongoing debate regarding the distinction between N1 and MMN by further suggesting that these reflect different hierarchical processing stages.

Two interesting research avenues can be derived from this study. First, we have shown that interval size can be a good predictor of N1m amplitude. However, a mechanistic link between this metric and neural responses is yet to be made. This would imply creating and refining a detailed computational model that links the acoustic properties of successive sounds with the corresponding neural activity. Second, our study shows how early sensory processing and higher-order probabilistic prediction can be disentangled. Therefore, experimental designs similar to ours might be a good way to test theories of cortical function such as predictive coding (Friston, 2005; Rao & Ballard, 1999) and active inference (Friston, FitzGerald, Rigoli, Schwartenbeck, & Pezzulo, 2017; Friston, Rosch, et al., 2017), which have hierarchical processing at their core.

## Conclusion

In this study, we aimed to determine whether the modulation of N1m amplitude by auditory surprise was different between musicians and non-musicians. Using a computational model of auditory expectation, we showed no differences between the groups in the otherwise clear association between neural responses and estimated surprise. Further exploratory analyses suggested that interval size and stimulus-specific adaptation, rather than probabilistic prediction, underlie the observed effects. This offers an explanation to why no effect of expertise was found, since we would expect higher-order probabilistic prediction, instead of early sensory processing, to be affected by the accuracy of musical knowledge. Interestingly, our results also suggest that auditory regularities and probabilistic prediction are reflected in later processing stages indexed by the MMNm and the P3am. Overall, our findings reveal a hierarchy of expectations in the auditory system, including early sensory processing, the formation of low-level regularities and higher-order probabilistic prediction. Therefore, our study constitutes an advance towards the understanding of hierarchical predictive processing in complex and realistic auditory contexts.

## Abbreviations

AIC: Akaike information criterion
BEM: Boundary element method
BF: Bayes Factor
ERF: Event related field
F_0_: Fundamental frequency
GMSI: Goldsmiths Musical Sophistication Index
IC: Information content
IDyOM: Information Dynamics of Music
MA: Mean amplitude
MET: Musical ear test
MNI: Montreal Neurological Institute
SSA: Stimulus specific adaptation
WAIC: Widely applicable information criterion

## Acknowledgments

We wish to thank the project initiation group, namely Christopher Bailey, Torben Lund and Dora Grauballe, for their help with setting up the experiments. We also thank Nader Sedghi, Massimo Lumaca, Giulia Donati, Ulrika Varankaite, Giulio Carraturo, Riccardo Proietti, and Claudia Iorio for assistance during MEG recordings. We are indebted as well to the group of Italian trainees fromIISS Simone-Morea, Conversano, who helped with the behavioral experiment. Finally, we thank Hella Kastbjerg for checking the English language of this manuscript. The Center for Music in the Brain is funded by the Danish National Research Foundation (DNRF 117), which did not have any influence on the scientific content of this article.

## Open practices

Materials and data for the study are available at https://osf.io/my6te/; DOI: 10.17605/OSF.IO/MY6TE

## CRediT authorship contribution statement

David R. Quiroga-Martinez: Conceptualization, Methodology, Software, Formal analysis, Data curation, Writing - original draft, Visualization, Investigation. Niels C. Hansen: Conceptualization, Methodology, Writing - original draft. Andreas Højlund: Conceptualization, Methodology, Software, Writing - original draft, Supervision. Marcus T. Pearce: Software, Formal analysis, Writing - original draft. Elvira Brattico: Conceptualization, Supervision, Writing - original draft. Peter Vuust: Conceptualization, Methodology, Supervision, Writing - original draft.

## Footnotes

1 Henceforth, an “m” will be added when referring to the magnetic counterpart of ERP components. When the “m” is omitted, we refer to the components in a more general sense, encompassing both their EEG and MEG manifestations.

