## Supplementary file 1 for "Decomposing neural responses to melodic surprise in musicians and non-musicians: evidence for a hierarchy of predictions in the auditory system"

Number and percentage of musicians playing a given instrument

| instrument | count | percentage |
|---|---|---|
| piano | 8 | 30.77 |
| voice | 4 | 15.38 |
| guitar | 3 | 11.54 |
| trumpet | 3 | 11.54 |
| bass | 2 | 7.69 |
| drums | 2 | 7.69 |
| cello | 1 | 3.85 |
| percussion | 1 | 3.85 |
| setar | 1 | 3.85 |
| violin | 1 | 3.85 |
