## Supplementary figures and images for "Decomposing neural responses to melodic surprise in musicians and non-musicians: evidence for a hierarchy of predictions in the auditory system"

### Supplementary file 2

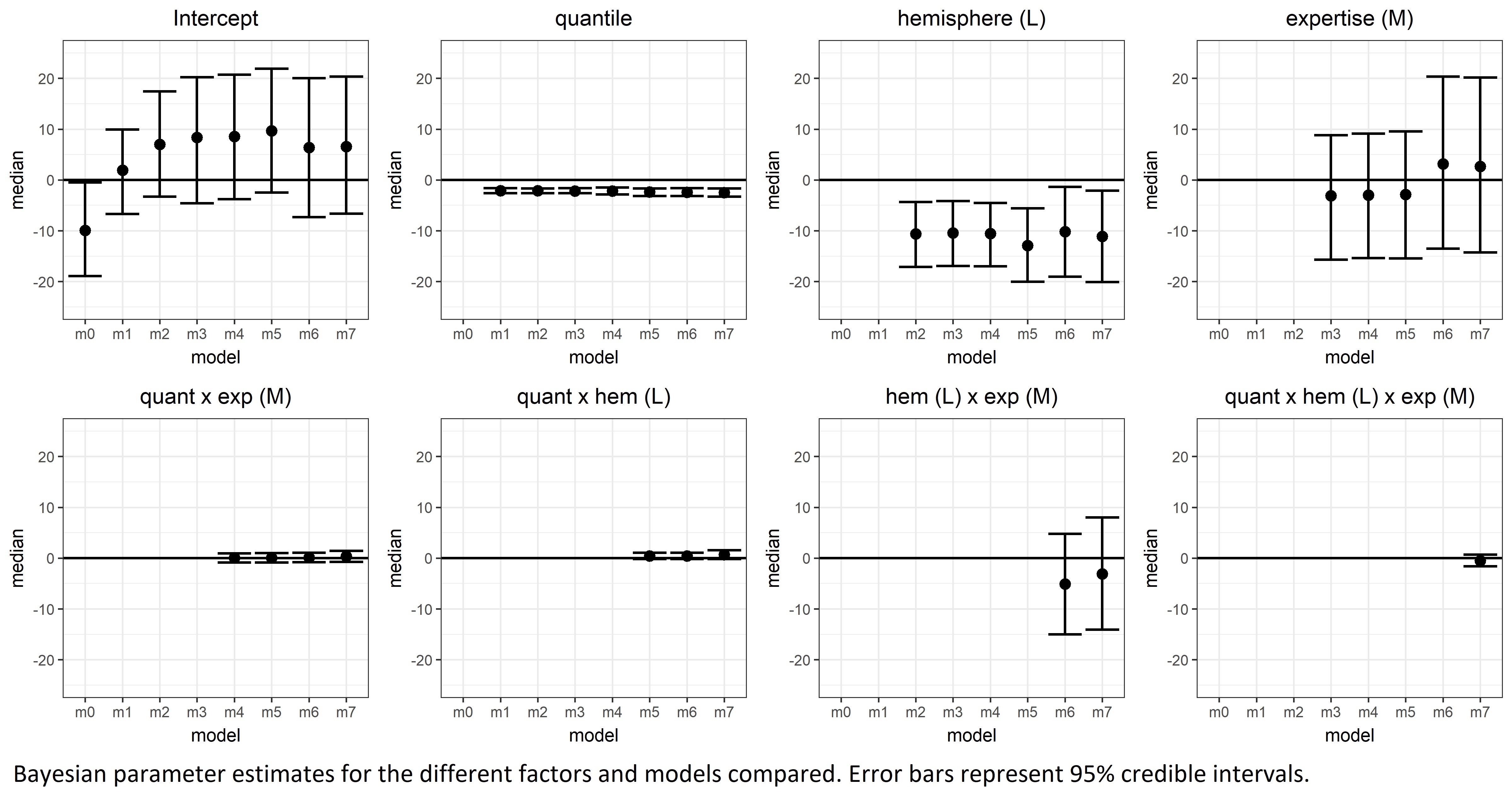
