## Supplementary file 3 for "Decomposing neural responses to melodic surprise in musicians and non-musicians: evidence for a hierarchy of predictions in the auditory system"

Acoustic similarity analysis of every possible pair of tones in our stimulus set. Cosine similarity (applied on the spectrograms) is used as a measure of how similar two acoustic waves are. Here we show that similarity decreases with pitch height, which is consistent with a sensory adaptation explanation of the effect of pitch interval size on the N1m response.

A) Power spectral density of two different tones.

Dashed vertical lines indicate their fundamental frequencies

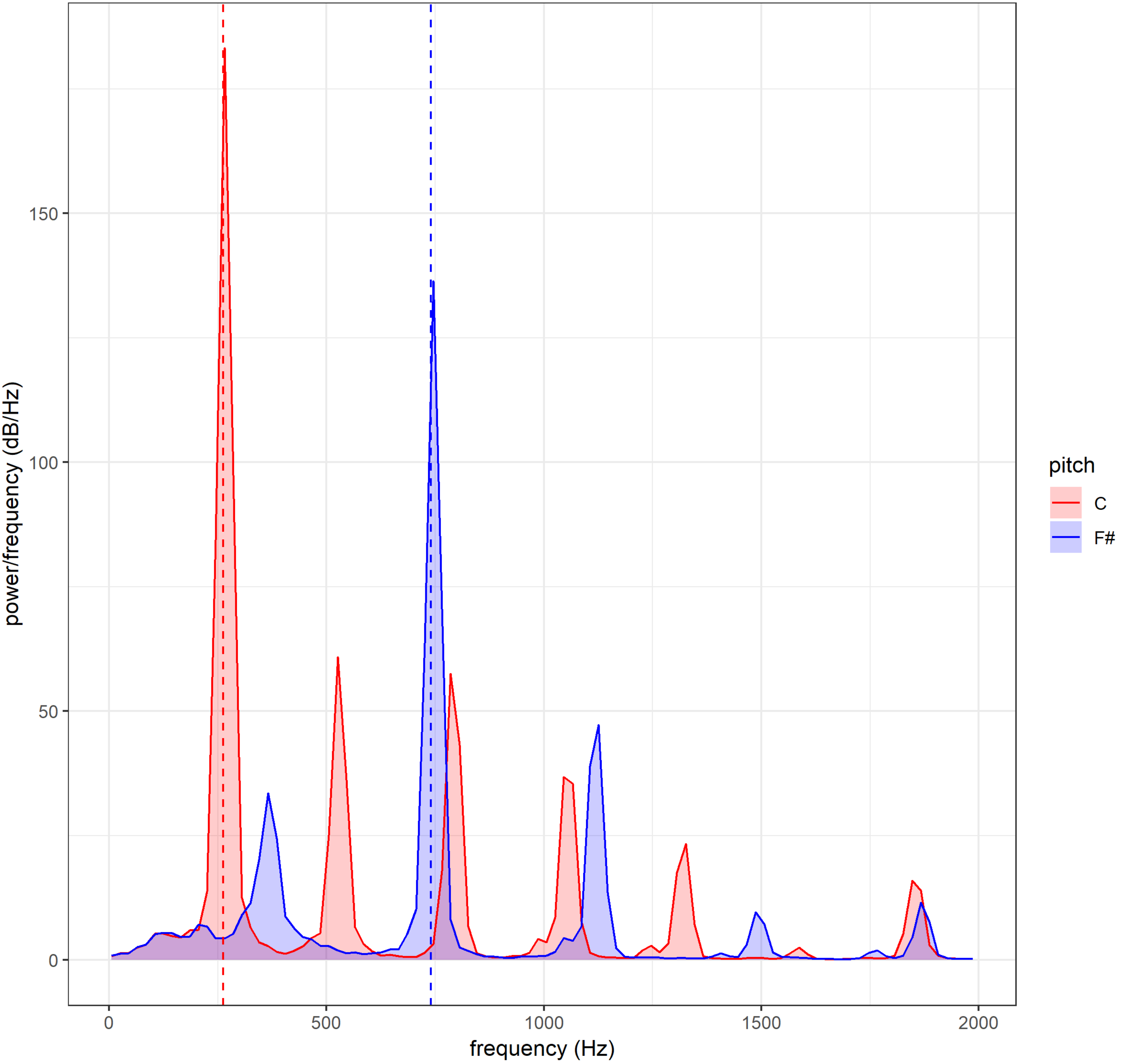

B) Cosine similarity matrix for every possible interval (from pitch 1 to pitch 2) in our stimulus set. Pitch identity is given in MIDI notation, where C4 = 60.

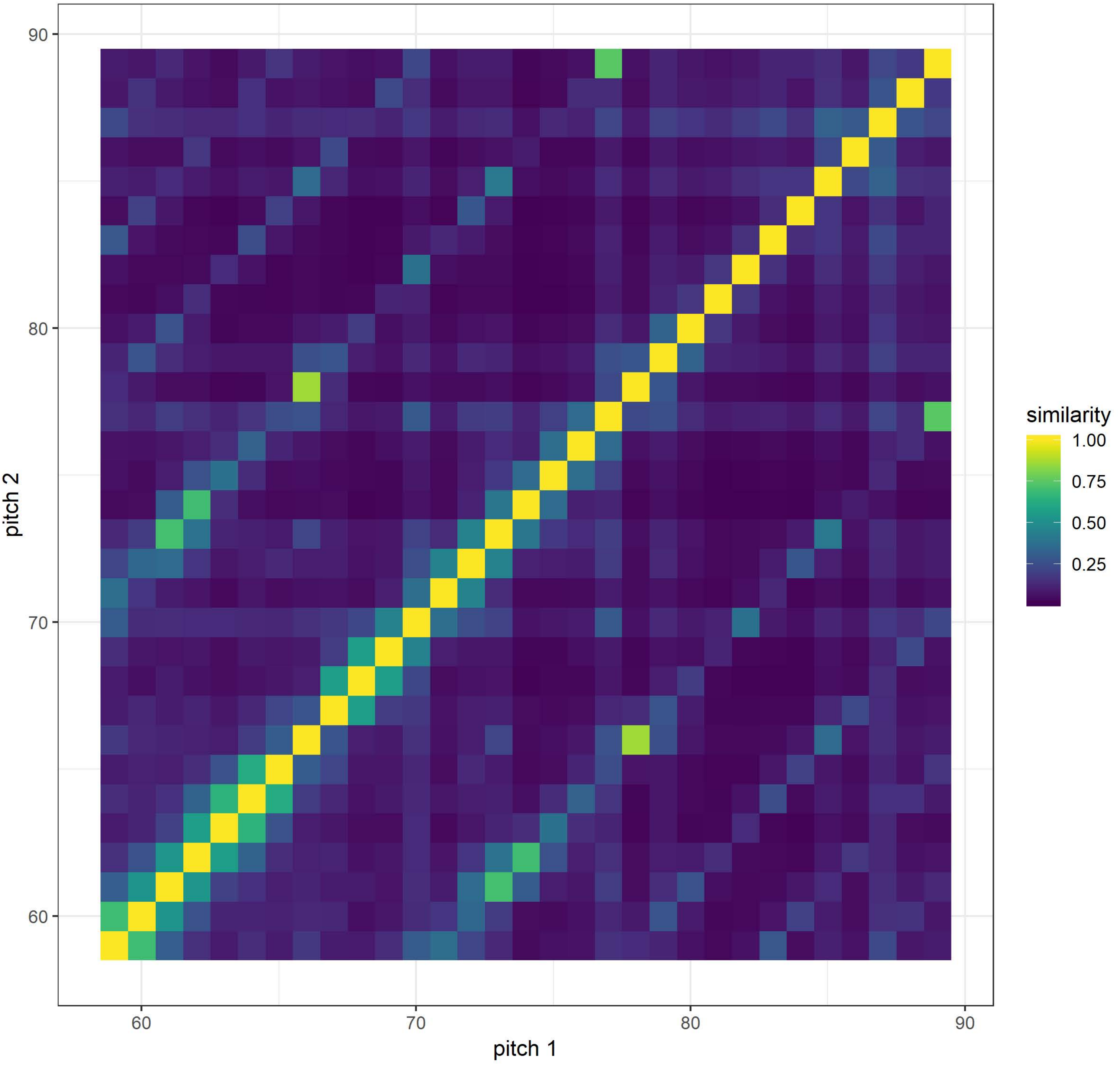

C) Spectral (cosine) similarity profiles for a low-pitched and a high-pitched tone

Pitch identity is given in MIDI notation, where C4 = 60. Note how similarity decreases faster for the higher pitch (C6 = 84).

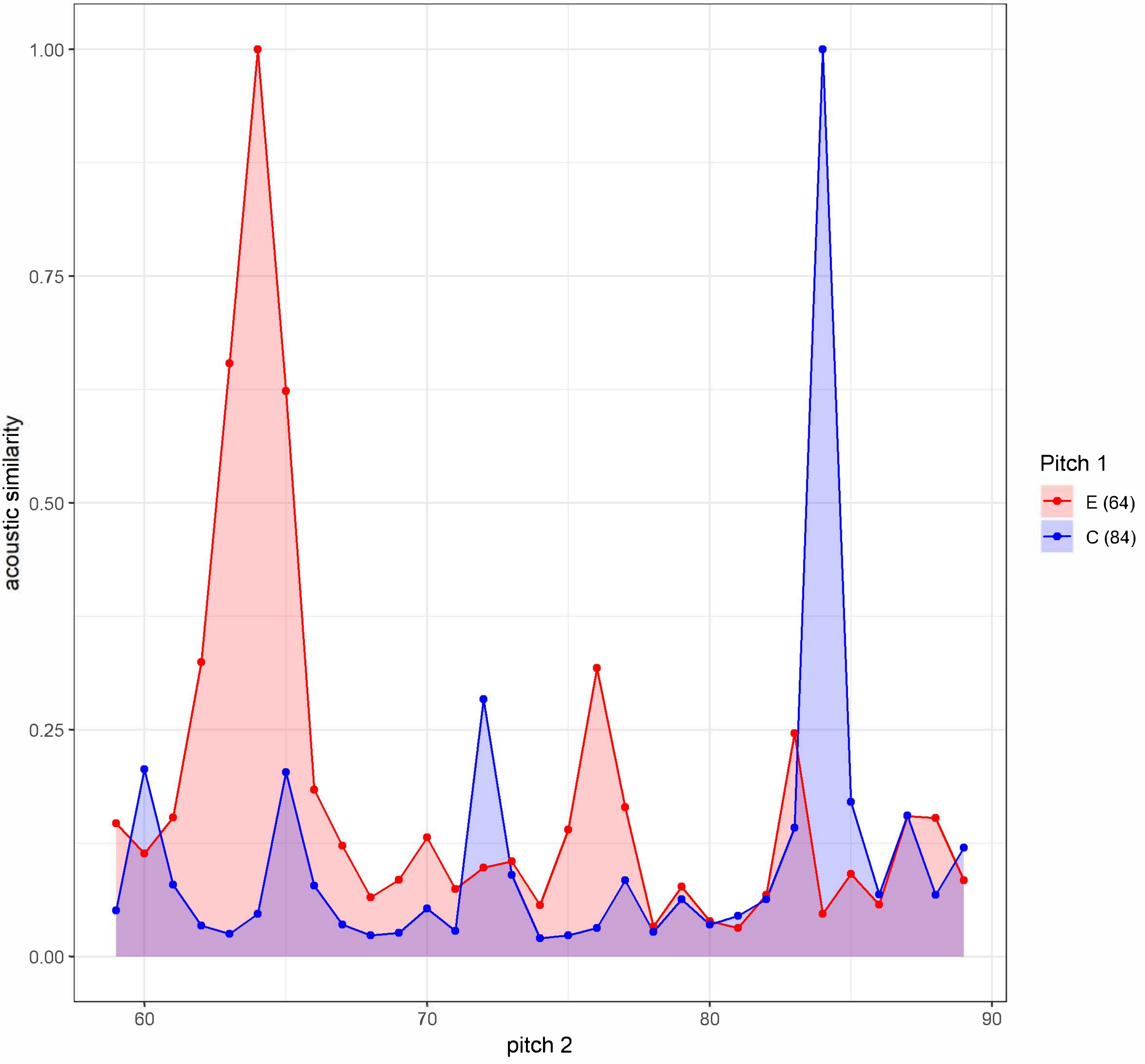

D) Mean spectral (cosine) similarity for each possible absolute interval in our simulus set.

Note how tones that are 12, 19, 24 and 28 semitones apart are more similar than it would be expected from pure pitch height. These correspond to octave, fifth and third musical intervals.

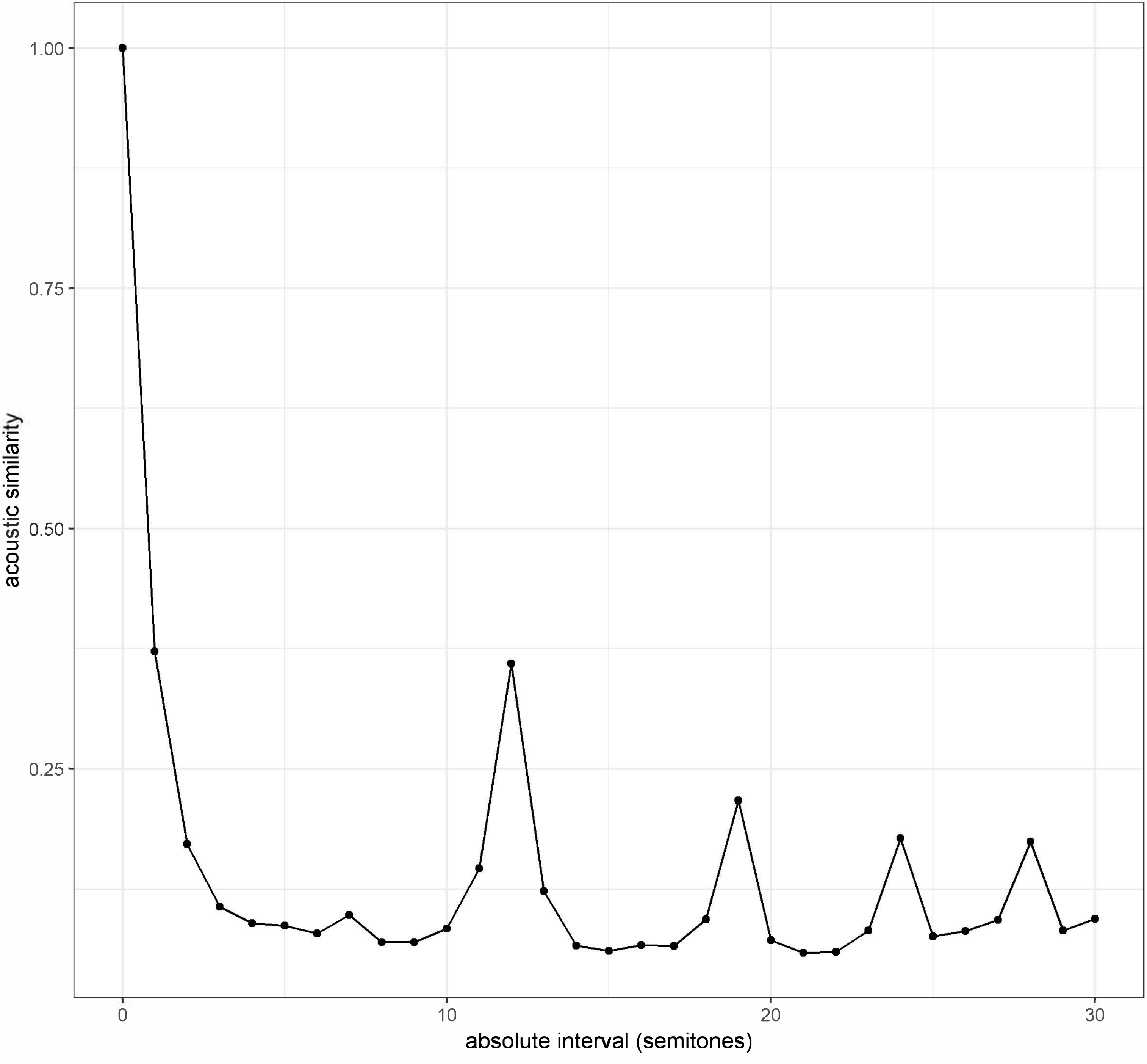
